# Rapid enhancement of antibiotic resistance and divergent remodeling of gene expression in single cells of bacteria

**DOI:** 10.64898/2026.10.22.760911

**Authors:** Miki Umetani, Mikihiro Hashimoto, Sayo Akiyoshi, Reiko Okura, Natsue Sakata, Hazuki Kotani, Edo Kussell, Chikara Furusawa, Yuichi Wakamoto

**Affiliations:** Department of Biology, New York University; Department of Basic Science, Graduate School of Arts and Sciences, The University of Tokyo; Center for Biosystems Dynamics Research, RIKEN; Department of Physics, New York University; Universal Biology Institute, The University of Tokyo; Research Center for Complex System Biology, The University of Tokyo

## Abstract

Exposing bacterial cell populations to antibiotics often results in the emergence of descendants with enhanced resistance. Phenotypic heterogeneity and global physiological responses are thought to contribute to resistance evolution. However, identifying the specific stages of drug exposure where phenotypic heterogeneity and physiological responses play critical roles remains elusive. Using long-term single-cell lineage tracking of an *Escherichia coli* resistance model, here we reveal that resistance-enhanced cells emerge rapidly within a few generations in a clonal bacterial cell population under chloram-phenicol (Cp) exposure and stably transmit the enhanced resistance for hundreds of generations in the presence of Cp. High expression of a resistance protein was advantageous during the initial period of drug exposure, but conferred only a marginal fitness advantage during the period of resistance enhancement. Transcriptomic profiles of resistance-enhanced cell populations were lineage-specific, suggesting that multiple gene expression states can support enhanced resistance. Altogether, these results suggest that an interplay between initial phenotypic heterogeneity and divergent remodeling of gene expression drives the rapid acquisition of enhanced antibiotic resistance.

## Main

The emergence and spread of antibiotic-resistant bacteria is a major health concern worldwide. Antibiotic resistance is a phenomenon in which bacterial cells grow and replicate in the presence of high concentrations of an antibiotic [1, 2]. Diverse resistance mechanisms to counter drug action have been documented, such as preventing the entry or accumulation of drugs, modifying drugs or drug targets, and bypassing metabolic pathways [3, 4]. Such remodeling of cellular functions is often caused by genetic changes inside cells, ranging from point mutations to the acquisition of novel genetic elements through horizontal gene transfer [3, 4]. Adaptive laboratory evolution and whole-genome sequencing have enabled the direct recording of resistance evolution and identified the transitions of genotype spectra in cellular populations [5–8].

However, genetic changes are not the sole driver of resistance evolution. Pre-existing non-genetic heterogeneity and physiological responses can also contribute to, or even be required for, adaptive evolution [9–15]. For example, stochastically produced or environmentally induced phenotypic heterogeneity in clonal bacterial populations allows a fraction of cells to survive lethal antibiotic exposure for extended periods [1]. Such non-genetic refractoriness, known as bacterial persistence, is thought to promote the evolution of antibiotic resistance [9–11, 14, 16]. Even sublethal concentrations of antibiotic drugs can induce physiological changes in bacterial cells, making them less susceptible to higher doses [17–20]. Similarly, *adaptive resistance*—a specific class of non-mutational resistance characterized by its transient nature—is well recognized [20]. In this mode of resistance, antibiotic exposure triggers various intracellular adaptive programs that make cells less susceptible to the drug, but cells revert to susceptible states when cultured in the absence of the drug [20–25]. However, a caveat is that at least some of the phenomena reported as adaptive resistance are plausibly driven mutationally by the selection of subpopulations with transient tandem DNA amplification—a frequently encountered form of unstable *heteroresistance* [26, 27].

Heterogeneous gene expression, such as that of toxin-antitoxin systems [28], efflux pumps [29], and drug-activating enzymes [30], has been characterized as a non-mutational factor correlated with individual cell survival under antibiotic exposure. A more straightforward example is the heterogeneous expression of proteins that predominantly confer resistance via direct drug inactivation or target modification [13]. In such cases, it is reasonable to assume that cells expressing higher levels of resistance-conferring proteins possess an adaptive advantage and are more likely to survive upon antibiotic exposure. However, antibiotics also provoke diverse physiological changes in cells [17, 19]. Thus, it remains unclear to what extent the heterogeneous expression of resistance-conferring proteins contributes to cell fate diversification. Furthermore, due to technical challenges in long-term single-cell lineage tracking during resistance evolution, it also remains elusive how rapidly resistance-enhanced cells emerge in a cell population under antibiotic exposure.

To address these questions, we constructed an *E. coli* model strain expressing a fluorescently tagged chloramphenicol resistance protein and tracked the dynamics of individual cells for hundreds of generations under various antibiotic exposure histories using custom microfluidic devices.

### Rapid acquisition and stable transmission of enhanced antibiotic resistance at the single-cell level under antibiotic exposure

To investigate single-cell responses and the enhancement of antibiotic resistance, we constructed an *E. coli* model strain, F3-VCAT/pmCh (Fig. 1A). This strain expresses a chloramphenicol acetyltransferase tagged with a yellow fluorescent protein (Venus-CAT) driven by a constitutive promoter (P_LtetO_*_−_*_1_ [31]) on the chromosome, as well as a red fluorescent protein (mCherry) also driven by the P_LtetO_*_−_*_1_ promoter from a low-copy plasmid. The Venus-CAT confers resistance to chloramphenicol (Cp), an antibiotic that inhibits protein translation by binding to the 50S subunit of the bacterial 70S ribosome, whereas the mCherry serves to visualize individual cell bodies during time-lapse microscopy.

**Figure 1:**
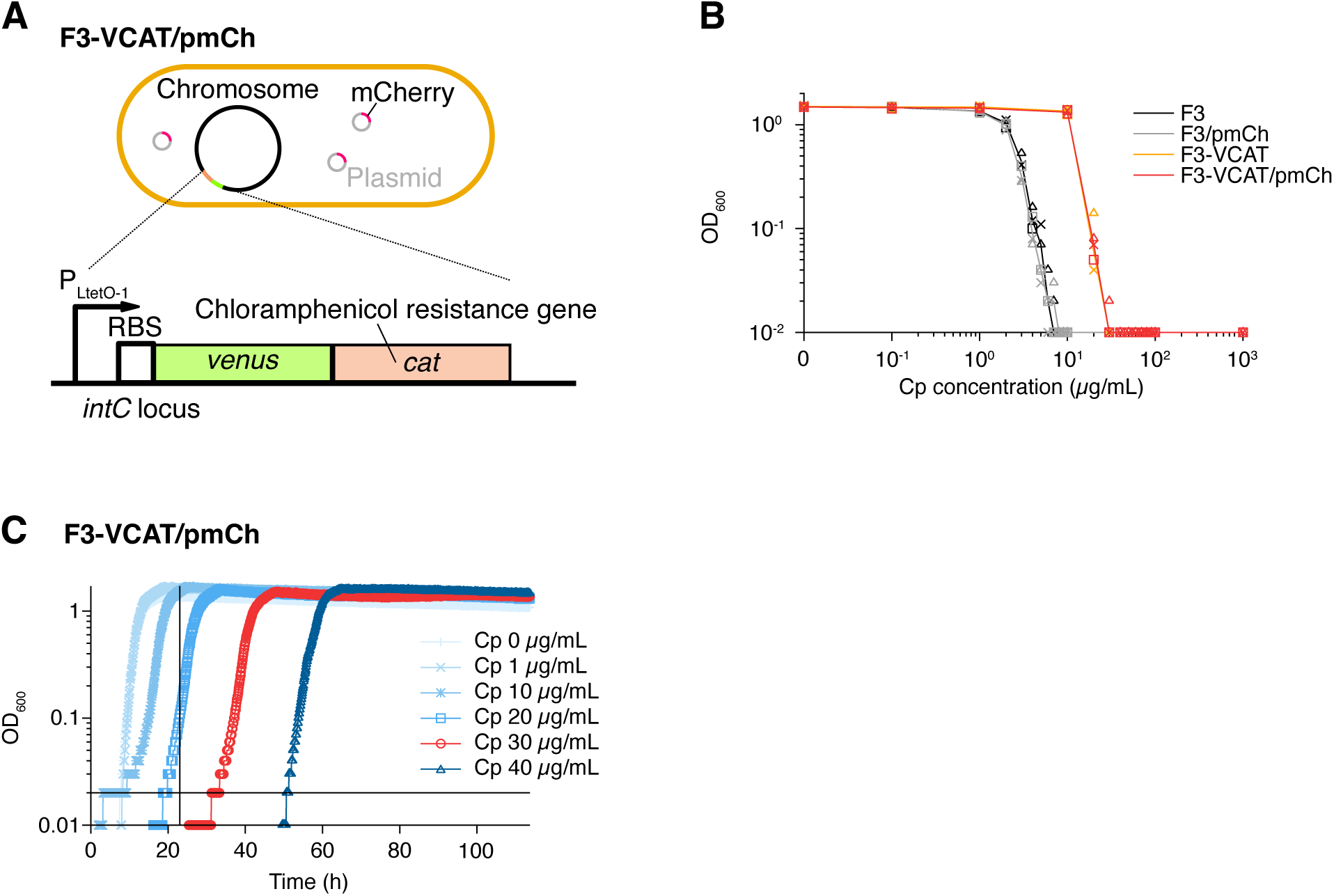
A synthetic *E. coli* model strain for studying antibiotic resistance enhancement. **A**. Schematic drawing of F3-VCAT/pmCh. This *E. coli* model strain constitutively expresses Venus-tagged chloramphenicol (Cp) acetyltransferase (Venus-CAT) from the *intC* locus on the chromosome under the control of the P_LtetO_*_−_*_1_ promoter [31]. A ribosome binding site (RBS) is positioned between the promoter and the Venus-CAT coding sequence. In addition, this strain carries a low-copy-number plasmid that expresses the red fluorescent protein mCherry, also under the control of the same promoter. **B**. MIC measurements against Cp. The plot shows the optical density at 600 nm (OD_600_) of *E. coli* cell cultures after 23 hours of incubation at 37°C in M9 medium containing different concentrations of chloramphenicol (Cp; horizontal axis). Black, gray, orange, and red represent the F3, F3/pmCh, F3-VCAT, and F3-VCAT/pmCh strains, respectively (“-VCAT” and “/pmCh” in the strain names indicate the presence of the *venus-cat* gene and mCherry-expressing plasmid, respectively). Data points (triangles, squares and crosses) indicate results for three biological replicates. Lines represent the geometric means of the replicates. Optical density values below 0.01 were plotted as 0.01, as such low readings are indistinguishable from background signal fluctuations. **C**. Population growth curves of F3-VCAT/pmCh under Cp exposure. Each series represents the growth of F3-VCAT/pmCh strain cultured in M9 medium containing 0, 1, 10, 20, 30, or 40 µg/mL of Cp. Measurements were performed simultaneously with *n* = 1. The initial cell density of the cultures was adjusted to an OD_600_ of 0.001. Only data points with OD_600_ *≥* 0.01 are shown, as spectrophotometer readings below this threshold are unreliable. The vertical line marks the 23-hour time point at which OD_600_ values were measured in the MIC test shown in B, and the horizontal line represents the OD_600_ threshold used for MIC determination. The red growth curve corresponds to the condition with 30 µg/mL Cp, the MIC for this strain.

Due to the presence of Venus-CAT, the Cp resistance of this model strain was higher than that of the control strain lacking this module (F3/pmCh); the minimum inhibitory concentration (MIC) of Cp was 30 µg/mL for F3-VCAT/pmCh, whereas it was below 10 µg/mL for F3/pmCh (Fig. 1B). The presence of the mCherry-expressing low-copy plasmid (pmCh) did not affect Cp resistance (Fig. 1B). Therefore, the increased resistance of the model strain was attributable to the Venus-CAT module.

Although the F3-VCAT/pmCh strain exhibited higher resistance, it remained susceptible to Cp at concentrations of 30 µg/mL (MIC) or higher, as a 23-h incubation in the standard MIC measurement did not show any detectable growth (Fig. 1B). However, when populations of this strain were incubated in the presence of Cp for longer than 23 h, regrowth and subsequent rapid exponential growth were observed, with delays that lengthened at higher Cp concentrations (Fig. 1C). These dynamics suggest that at least a fraction of cells in the population acquired enhanced resistance, enabling them to grow in the presence of high Cp concentrations.

Because batch-culture-based assays cannot resolve the fine details of heterogeneous single-cell dynamics under Cp exposure, we performed single-cell time-lapse measurements using a membrane-covered microchamber array (MCMA) device [32]. In this setup, *E. coli* cells were enclosed in microchambers microfabricated on a glass coverslip and sealed with a semipermeable cellulose membrane [32]. Cells grow in a monolayer, and the surrounding medium conditions can be maintained or dynamically shifted at arbitrary times by controlling the medium flow above the membrane.

When growing microcolonies of F3-VCAT/pmCh were exposed to 30 µg/mL Cp (the MIC, Fig. 1B, C), the growth and division of all individual cell lineages were suppressed (Fig. 2A, S1, S2, Movie S1 and S2). Although most cells eventually ceased growing and dividing (green lineage in Fig. 2A-C; pink lineage in Fig. 2D, E; and Fig. S1 and S2), a subset of cell lineages continued to grow and divide at reduced rates (orange lineage in Fig. 2A-C; yellow lineage in Fig. 2D, E; Fig. S1, Movie S1, and S2). Some growth-arrested cells exhibited marked shrinkage of their cell bodies after arrest, suggesting a loss of viability (Fig. S2), whereas others showed little or no such morphological change (Fig. 2A and S1). Importantly, within just a few cell divisions under Cp exposure—during which many sibling cells entered growth arrest—the resistance became stable, with nearly all descendant cells continuing to grow and divide without arrest in the presence of Cp (Fig. 2B, D, Movie S1 and S2). Single-cell tracking revealed that 12% of cells at the onset of Cp exposure (25/208) yielded lineages with such stably growing, resistance-enhanced descendant cells.

**Figure 2:**
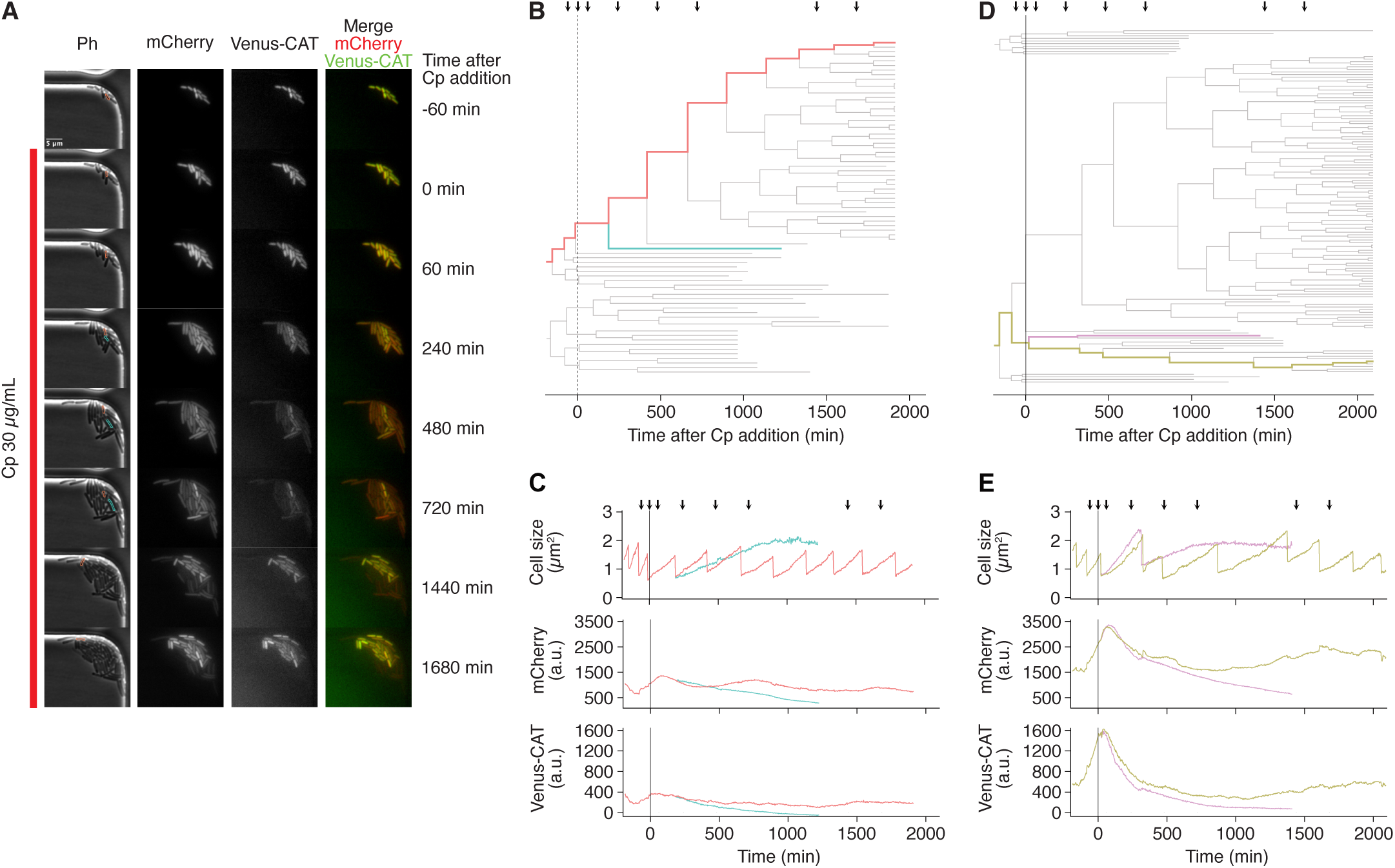
Heterogeneous response of individual cells to Cp exposure at MIC. **A**. Time-lapse images of representative F3-VCAT/pmCh cells exposed to 30 µg/mL Cp in the MCMA device. The images show, from left to right: phase-contrast, mCherry fluorescence, Venus-CAT fluorescence, and merged images of mCherry (red) and Venus-CAT (green) channels. The red line to the left of the images indicates the period of Cp exposure. A representative growing cell in this environment is outlined in orange, and a representative growth-arrested cell is outlined in green. These two cells are sister cells derived from the same division event after the onset of Cp exposure. **B**. Single-cell lineage trees corresponding to the microcolony shown in panel A. Each bifurcation represents a cell division. Branches that terminate before the final time point indicate that tracking of growth-arrested cells became infeasible due to significant loss of mCherry fluorescent signal. Colored lineages correspond to the cells highlighted in the same colors in panel A. The vertical line indicates the time point of Cp exposure onset (*t* = 0). Black arrows indicate the time points at which the micrographs in panel A were taken. **C**. Time series of cell size (top), mCherry fluorescence intensity (middle), and Venus-CAT fluorescence intensity (bottom) in single-cell lineages. Fluorescence intensity is calculated as the total signal within the detected cell boundary divided by the cell area, serving as a proxy for the intracellular concentration of each protein species. The orange and green traces correspond to the cell lineages highlighted in the same colors in panels A and B. **D**. Lineage trees corresponding to the microcolony shown in Fig. S1, with colored branches corresponding to the cells highlighted in yellow and pink. The vertical line indicates the time point of Cp exposure onset (*t* = 0). Black arrows mark the time points of the images in Fig. S1. **E**. Time series of cell size (top), mCherry fluorescence (middle), and Venus-CAT fluorescence (bottom) in the two highlighted lineages in panel D and Fig. S1.

### Long-term single-cell tracking reveals the stability of enhanced resistance under Cp exposure and its rapid reversion to a susceptible state upon Cp removal

To confirm the stability of the enhanced resistance in growth-continuing cells over extended generations, we performed long-term single-cell lineage tracking using a different microfluidic device, the “dynamics cytometer” [33]. This device allows the tracking of single cells trapped in narrow growth channels by discarding excess cells into wider flow channels. The environmental conditions surrounding the cells are controlled by medium exchange across a semipermeable membrane, as in the MCMA device [32, 33].

The initial responses of cells exposed to 30 µg/mL Cp in the dynamics cytometer were consistent with those observed in the MCMA device; Cp exposure suppressed the growth of most cell lineages (Fig. 3A, B, and Movie S3), but a small fraction of cells continued growth and division (Fig. 3B and Movie S3). Extended observation revealed that the progeny of these growing cells indeed stably transmitted enhanced resistance and produced almost no growth-arrested cells over 100 generations (for more than two weeks) under a continuous flow of 30 µg/mL Cp (Fig. 3B and Movie S3).

**Figure 3:**
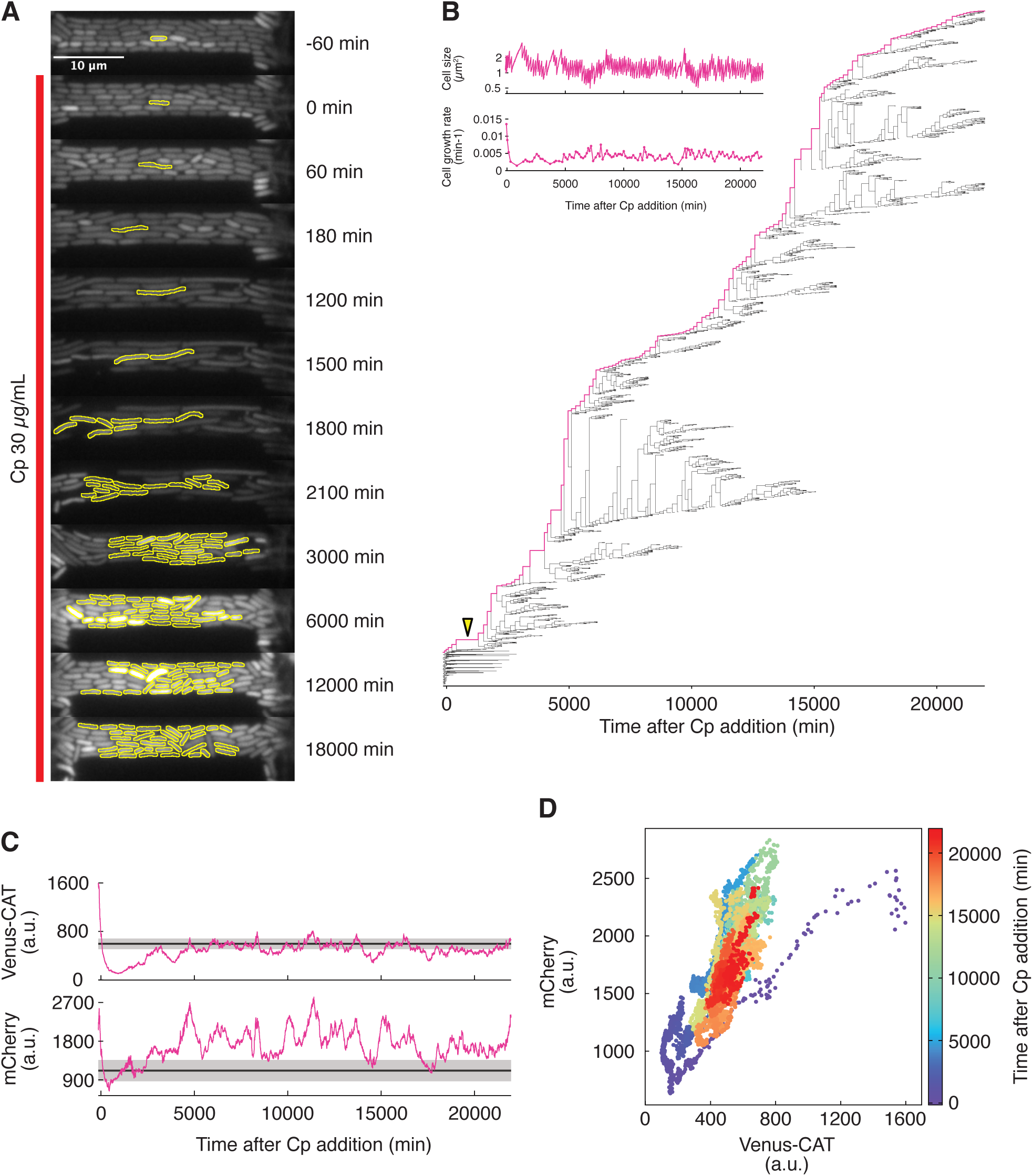
Long-term inheritance of enhanced resistance under continuous Cp exposure. **A**. Time-lapse mCherry fluorescence images of F3-VCAT/pmCh cells exposed to 30 µg/mL Cp in the dynamics cytometer. Cells outlined in yellow represent a cell lineage that inherited enhanced resistance and continued to divide for more than 100 generations under Cp exposure. The red line to the left indicates the period of Cp exposure. **B**. Single-cell lineage trees corresponding to the cells shown in panel A. Each bifurcation represents a cell division. The lineage highlighted in pink corresponds to a representative cell that remained in the growth channel throughout the observation. The inset shows temporal changes in cell size and growth rate along this pink lineage. **C**. Temporal changes in Venus-CAT and mCherry fluorescence levels. The bold solid lines indicate the mean values among cells present at the time of drug addition, and the shaded areas represent the standard deviations. **D**. Correlation between Venus-CAT and mCherry fluorescence levels. The color scale indicates time after Cp addition.

This long-term single-cell tracking characterized the slow recovery dynamics of Venus-CAT and mCherry expression. The expression levels of Venus-CAT significantly decreased in response to Cp exposure but gradually recovered to levels comparable to the pre-exposure population mean, taking approximately 5,000 min (3.5 days) (Fig. 3C). The expression levels of Venus-CAT and mCherry in single cells were tightly correlated (Fig. 3D). Consequently, similar recovery dynamics was also observed in mCherry expression levels, although its basal expression levels after recovery were higher than those before Cp exposure (Fig. 3C).

This single-cell tracking setup provided additional evidence for the enhanced resistance of cells growing under Cp exposure. When the concentration of Cp was increased to 40 µg/mL for cells that were growing and dividing stably in the presence of 30 µg/mL Cp, most of them continued growing and dividing, albeit at a reduced rate (Movie S4 and Fig. S3). However, when cells were exposed to 40 µg/mL from the beginning, all cells in the growth channels immediately stopped dividing and eventually halted growth (Movie S4 and Fig. S3). Therefore, cells stably growing at the MIC can resist an even higher concentration that would otherwise prohibit the vast majority of cells from continuing growth. We confirmed that the resistance permitting stable growth and division can be enhanced up to at least 60 µg/mL through a stepwise increase in Cp concentration in the dynamics cytometer (Movie S5).

In addition, single-cell measurements confirmed the transient nature of this enhanced resistance. When Cp was temporarily removed for 8.1 h (ca. 8–9 generations) from cells growing stably in 30 µg/mL Cp, the cells lost their resistance: no continuously growing cells appeared among their progeny upon a second exposure to 30 µg/mL Cp (Movie S6). Therefore, the enhanced resistance acquired under Cp exposure can be rapidly lost, at least in some cell lineages, once Cp is removed from the environment.

### The expression level of the resistance protein in individual cells plays a diminishing role in fitness under prolonged Cp exposure

Because only a subset of cell lineages continued to grow and acquired enhanced resistance, we next investigated the factors determining these distinct cellular fates under Cp exposure. Given that this model strain expresses Venus-CAT, the expression level of this resistance protein in individual cells was a primary candidate that contributed to enhanced resistance.

The inspection of Venus-CAT profiles along individual cell lineages measured in the MCMA device identified both high- and low-Venus-CAT-expressing cells giving rise to growth-continuing cell lineages upon Cp exposure (Fig. 2C and E). To quantitatively evaluate the contribution of Venus-CAT expression to cell fate diversification, we assessed the correlations between its expression level and the fitness of both cell populations and individual cells, using the data collected in the MCMA device. To this end, we first identified cell lineage branches that lost division ability using a Bayesian inference method based on the traits of growth-continuing and growth-arrested cells (Fig. S4; see Materials and Methods for details). By excluding these non-dividing branches from single-cell pedigree trees, we characterized the population dynamics under Cp exposure in the MCMA device (Fig. 4A), the dynamics of phenotypic traits in individual cells (Fig. 4B), and the number of viable descendant cells originating from each individual cell. Here, we define a “viable cell” as one that retains division ability; although Cp is generally classified as a bacteriostatic antibiotic, our time-lapse observations showed that cells remaining non-dividing for extended periods ultimately undergo cell death (Fig. S2).

**Figure 4:**
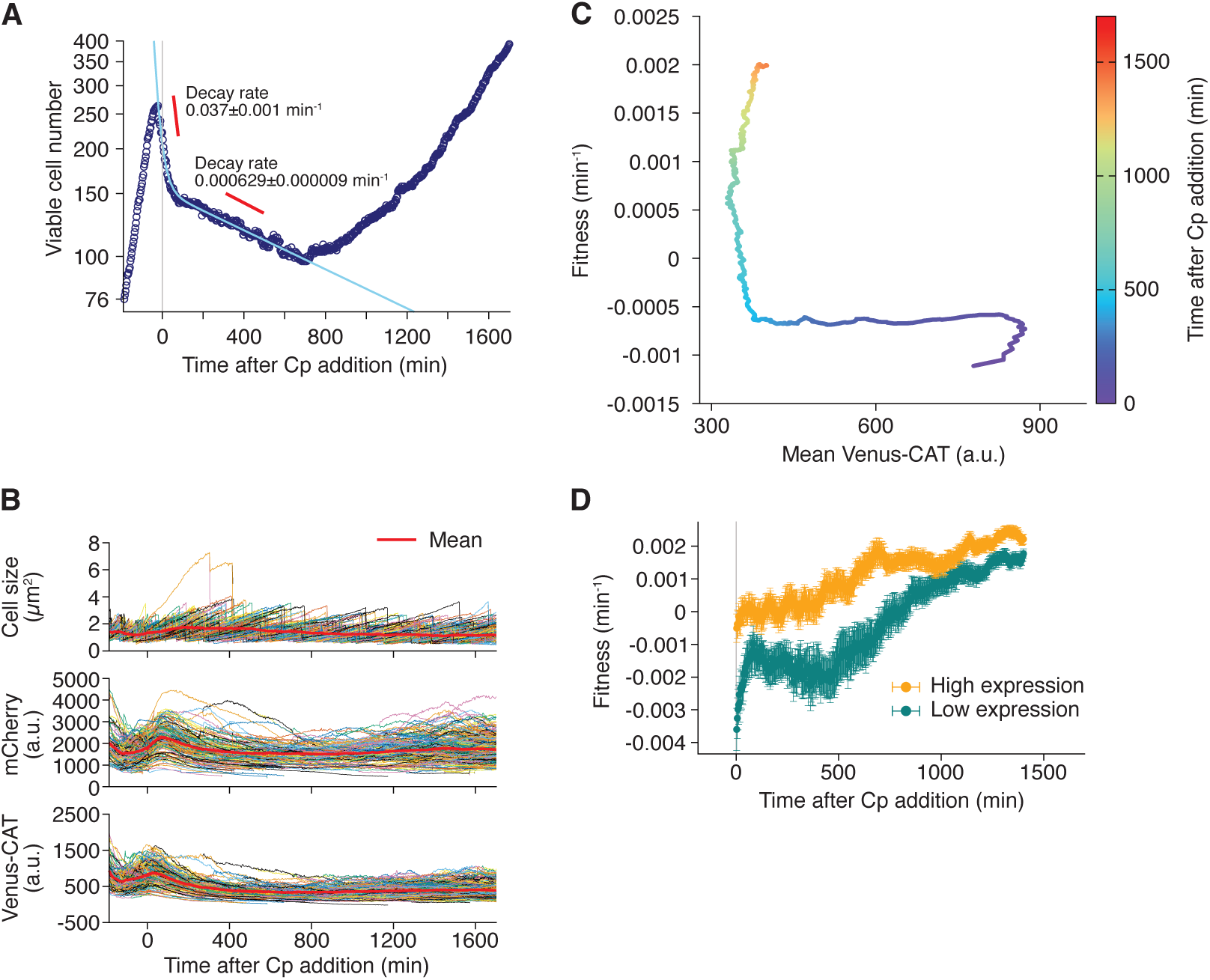
Limited contribution of resistance protein expression to fitness in the regrowing phase under Cp exposure. **A**. Time series of viable cell numbers in the presence of 30 µg/mL Cp (MIC) in the MCMA device. Navy points represent the logarithm of viable cell numbers counted in the MCMA device at each time point. The light blue line represents a fit of double exponential decay curve to the data in the decreasing phase: *N* (*t*) = 54.1 exp(*−*3.68 *×* 10*^−^*^2^*t*) + 1.52 *×* 10^2^ exp(*−*6.29 *×* 10*^−^*^4^*t*). The gray vertical line indicates the time point of Cp addition. The number of viable cells started to decrease before Cp exposure because “viable cells” were defined as those capable of producing daughter cells; the branches of Cp-exposed cells that did not divide even once under Cp exposure were removed from cellular pedigree trees (see Materials and Methods for details). **B**. Time series of cell size (top), mCherry fluorescence intensity (middle), and Venus-CAT fluorescence intensity (bottom) in 1,243 viable single-cell lineages. The bold red line in each graph indicates the mean value at each time point. **C**. Correlation between mean Venus-CAT fluorescence level and fitness after MIC-level Cp exposure. Colors indicate the time after Cp addition. The mean Venus-CAT expression level corresponds to the bold red line in the bottom panel of B. Fitness was calculated as the slope of the viable cell number dynamics (panel A) using a 300-minute time window. **D**. Comparison of fitness between high and low Venus-CAT expression groups. At each time point, cells were classified into the top 50% (orange) and bottom 50% (green) based on their Venus-CAT expression levels. To calculate the fitness of each group, we counted the number of viable progeny cells, *N_p_*, at a 300-min later time point that originated from the ancestor cells in each group. Fitness was then calculated as (1*/*Δ*t*) ln*{N_p_/N}*, where *N* is the number of ancestor cells in each group and Δ*t* = 300 min is the time window. Error bars represent standard deviations calculated from 10,000 bootstrap resamples, in which single-cell lineages in each group were randomly selected.

The population dynamics constructed from multiple single-cell pedigree trees originating from 76 ancestral cells revealed an initial biphasic decline in viable cell numbers upon exposure to Cp at the MIC (Fig. 4A). These dynamics mirror those observed in antibiotic persistence, although persistence is traditionally defined only in the context of bactericidal antibiotics [1]. Following this biphasic decay, the number of viable cells in the population began to increase steadily at approximately 700 min after the onset of Cp exposure, which aligns with the afore-mentioned observation that growth-continuing cells stably transmitted enhanced resistance to nearly all their descendant cells (Fig. 2B and D).

We evaluated the slope of the logarithm of the population dynamics as the population fitness at each time point, using a 300-min time window for slope calculation. Then, we investigated how these fitness transitions were correlated with the Venus-CAT expression level of the population. This analysis revealed that the increase in population fitness during the regrowing phase was not accompanied by a recovery of the expression level of the Venus-CAT protein (Fig. 4C), implying the contribution of additional factors to the resistance enhancement.

To further characterize the contribution of Venus-CAT expression to fitness, we classified individual cells at each time point into high-and low-expression groups (above and below the median, respectively) based on their Venus-CAT expression levels. We then counted the number of viable descendant cells 300 min later for each group to quantify its fitness from the fold change in viable cell numbers. We thereby found that while the fitness of the high-expression group was higher than that of the low-expression group during the initial phase of Cp exposure, this difference diminished under prolonged exposure (Fig. 4D). Importantly, the fitness of both groups recovered during later stages despite Venus-CAT expression levels remaining low (Fig. 4C and D). These results demonstrate that high expression of the resistance protein conferred a fitness advantage on individual cells during early phases of Cp exposure, but that the subsequent resistance enhancement was primarily driven by factors other than the resistance protein.

### Resistance-enhanced populations from different ancestral cells display distinct transcriptomic profiles

To identify the factors underlying the enhanced resistance, we next attempted to isolate cell populations originating from individual resistance-enhanced ancestral cells under Cp exposure. To this end, we seeded individual cells in 96-well plates by limiting dilution (the expected number of cells per well was 0.78; Fig. S5A) and incubated them in medium containing 30 µg/mL Cp. Stable regrowth was detected in only 23*±*7% (*n* = 4) of the wells, which was significantly lower than the fraction (54%) observed without Cp. The regrowth of Cp-exposed cells was detected after prolonged and variable lag durations, ranging from 80 to 130 h, whereas the regrowth of unexposed cells was detected almost simultaneously across wells at around 20 h (Fig. S5B). These regrowth behaviors of Cp-exposed cells were consistent with our single-cell observations that only a fraction of cells acquires enhanced resistance after several divisions under Cp exposure (Fig. 2, 3, S1, Movie S1 and S2). We selected three of the stably regrowing lineages under Cp exposure (A3, E9, and H10) from the 96-well plates and subjected them to transcriptomic profiling to characterize their gene expression states.

The transcriptomic profiles measured with microarrays revealed markedly distinct gene expression patterns among the resistance-enhanced cell populations derived from different ancestral cells under Cp exposure (Fig. 5). Among the genes exhibiting the top 10 and bottom 10 fold changes in expression in A3, many are regulated by RcsAB (Fig. 5A). RcsAB is known to upregulate genes involved in capsule formation and downregulate those related to the flagellar machinery [34]. These expression changes were indeed observed in the transcriptomic profile of A3 (Fig. 5A). Consistent with this observation, A3 cells formed capsules around their cell bodies at 37*^◦^*C, a phenotype typically observed only at low temperatures in laboratory strains of *E. coli* (Fig. S7A and B) [35]. These prominent expression changes in A3 cells were absent in E9 and H10 cells (Fig. 5B and C). In contrast, top 10 genes exhibiting the largest fold changes in expression in E9 cells include many genes regulated by CysB (Fig. 5B). The CysB-regulated genes are involved in sulfur assimilation and cysteine biosynthesis, with additional suggested contributions to diverse functions, including antibiotic resistance [36]. However, the expression changes in these genes were absent or less pronounced in A3 and H10 cells, as well as in the bottom 10 genes in E9 (Fig. 5A and C). The H10 cells exhibited relatively minor changes in their transcriptomic profile compared with A3 and E9 (Fig. 5C), although this cell lineage also grew stably under Cp exposure at a rate comparable to or even higher than the other two (Fig. S6A and B). Indeed, the number of genes exhibiting a 10-fold or greater expression change was only one in H10, whereas there were 110 and 16 such genes in A3 and E9, respectively (Fig. 5D). The observed diverse transcriptomic profiles indicate that multiple gene expression states can underlie the enhanced resistance state.

**Figure 5:**
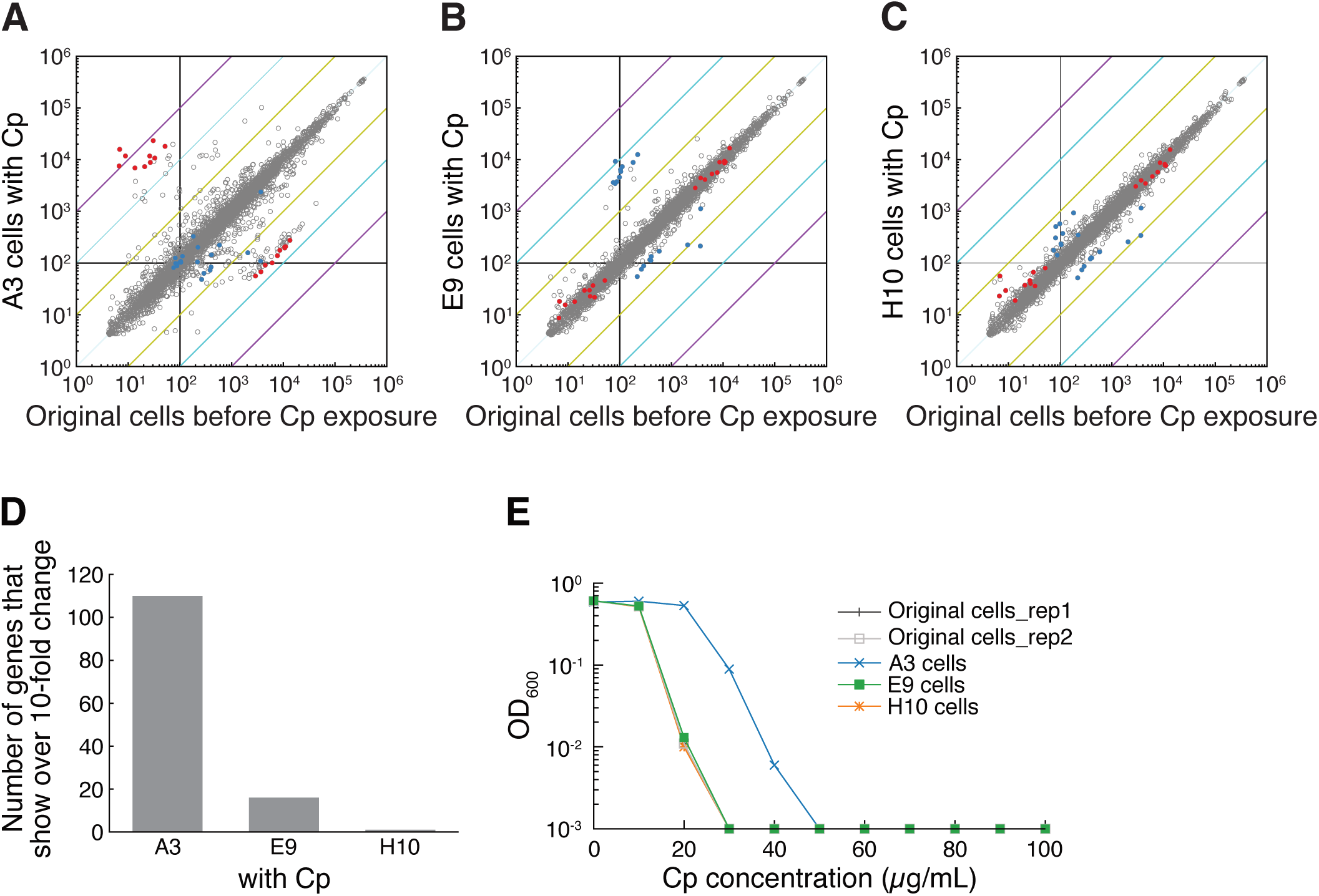
Distinct adaptive transcriptomic states of three cell lineages derived from different ancestral cells under Cp exposure. **A-C**. Scatter plots showing the transcriptome-wide comparison of gene expression levels between the original cell population prior to Cp exposure and each of the three resistance-enhanced cell lineages isolated in the presence of 30 µg/mL Cp (A: A3 cells, B: E9 cells, C: H10 cells). Each dot corresponds to a single gene. Genes exhibiting the top 10 and bottom 10 fold changes in expression in A3 and E9 under Cp treatment relative to the original cells before Cp exposure are highlighted in red and blue, respectively. Black lines represent the baseline threshold below which expression levels are considered too low for reliable interpretation. Yellow, cyan, and purple diagonal lines indicate 10-fold, 100-fold, and 1000-fold changes, respectively, providing visual references for the magnitude of expression shifts. **D**. The number of genes exhibiting more than 10-fold changes in transcript abundance in each resistance-enhanced cell lineage relative to the unexposed original cells. **E**. MIC measurements of the three resistance-enhanced cell lineages. The A3, E9 and H10 cell populations were passaged in Cp-free medium and re-exposed to a gradient of Cp concentrations to evaluate resistance stability. The unexposed original cell population was included as a reference.

To further characterize the distinctive properties of the isolated cell lineages, we removed Cp from the cultures of A3, E9, and H10 and grew them under Cp-free conditions for approximately 20 generations, including entry into stationary phase and pre-culture periods. We then subjected the exponentially growing cells to MIC measurements and transcriptomic profiling to assess any retained transcriptional or phenotypic memory in each lineage.

The MICs of E9 and H10 were identical to that of the parental unexposed cells (Fig. 5E). Therefore, these cell populations lost their enhanced resistance during the cultivation in the absence of Cp. However, A3 retained enhanced resistance even after Cp removal, with its MIC increasing from 30 to 50 µg/mL (Fig. 5E). The transcriptomic profiles after Cp removal mirrored these MIC results; the transcriptomic profiles of E9 and H10 returned to a state close to that of original Cp-unexposed cells, whereas A3 retained its profile observed under Cp exposure (Fig. S8).

Interestingly, when cells were incubated under Cp exposure for longer than 23 h (the duration used in standard MIC measurements), A3 and H10 cells exhibited delayed regrowth at high Cp concentrations where no regrowth was detected for the original unexposed cells or E9 (Fig. S6C-G). Although such resistance memory in H10 was not evident in its transcriptomic profile under Cp-free conditions (Fig. S8E and S8F), a fraction of cells within the H10 population might have retained enhanced resistance even after Cp removal.

To investigate whether genetic changes contributed to the observed resistance patterns in A3, E9, and H10, we subjected these lineages to whole-genome sequencing. Mutations were called using breseq [37] from MiSeq reads of these cell lineages—both before and after cultivation under Cp-free conditions—as well as their parental Cp-unexposed cells, followed by Sanger sequencing confirmation for ambiguous loci. However, we detected no genotypic differences between these resistance-enhanced cells and the original unexposed cells, even in A3 (Table S2). Although we cannot exclude the possibility of undetected genetic changes, such as tandem repeat DNA amplification, these results are consistent with resistance enhancement and transcriptomic modulation being driven primarily at the physiological level.

### Evaluating the correlation between resistance patterns and expression levels of selected resistance-associated genes

We next examined whether expression changes in candidate resistance-associated genes could account for the observed resistance patterns of the isolated cell lineages. First, these resistance patterns could not be explained by the expression levels of the Venus-CAT resistance protein; despite exhibiting a higher MIC, the expression level in A3 was nearly equivalent to that of the original unexposed cells (Fig. S9A). Although the mean expression level of Venus-CAT in H10 was slightly higher than that in the parental cells (Fig. S9A), this difference did not translate to an increase in MIC, as noted above (Fig. 5E).

Previous studies have identified genes implicated in Cp resistance [6, 18, 38–40]. Nevertheless, our transcriptomic analysis could not attribute the observed resistance phenotypes in these cell lineages to the expression patterns of those candidate genes (Fig. S10). For example, an evolutionary experiment using *E. coli* under dynamically adjusted Cp concentrations identified mutations in multiple genes in the evolved, resistance-enhanced cell populations [6]. However, expression changes in these genes in our isolated lineages did not consistently mirror their resistance patterns (Fig. S10A). The transcription factors Rob, MarA, and SoxS regulate the *mar-sox-rob* regulon, which includes efflux pumps and other components involved in antibiotic resistance [38, 39]. Yet, we did not detect pronounced expression changes in these transcription factors that paralleled the resistance profiles (Fig. S10A and B). Key resistance factors, such as *micF* and *ompF*, exhibited prominent expression changes in A3, but not in E9 or H10 (Fig. S10B). Additionally, *gadA* and *gadB*, which contribute to acid resistance [40] and potentially AcrAB-mediated multidrug efflux [18], were upregulated in all the cell lineages under Cp exposure, particularly in A3 (Fig. S10C). However, their expression levels remained elevated in E9 even under Cp-free conditions, which cannot account for its MIC being comparable to that of the original unexposed cells. Altogether, none of the single genes analyzed above can explain the resistance patterns across all cell lineages, although some may primarily determine the resistance level of a specific lineage.

Because the genes regulated by RcsAB and CysB exhibited pronounced expression changes in A3 and E9, respectively (Fig. 5A and B), we examined whether the induction of these transcriptional regulators alone could confer enhanced resistance. The IPTG-inducible expression of either RcsA or CysB revealed that overexpression of these regulators was insufficient to increase the MIC (Fig. S9B and C), even though RcsA induction resulted in capsule formation (Fig. S7C). In fact, for CysB, overexpression even resulted in a slight decrease in MIC (Fig. S9C). These results indicate that altering the expression of RcsA or CysB alone is insufficient to reproduce the enhanced resistance of A3 or E9, respectively, suggesting that additional physiological changes are required. The resistance phenotypes therefore appear to reflect broader cellular states rather than simple expression changes in genes governed by any single tested regulator.

### Absence of initial nucleoid compaction in growing cells upon Cp exposure

Transcriptomic profiling of the resistance-enhanced cells revealed lineage-specific expression states accompanied by expression changes in numerous genes (Fig. 5 and S8). Considering the impact of nucleoid structure on genome-wide transcription in bacterial cells [41], we visualized nucleoid structural changes in individual cells and investigated their relevance to growth continuation under Cp exposure.

To visualize nucleoids, we constructed an *E. coli* strain, F3-VCAT-HupAmCh. This strain expresses the nucleoid-associated protein HupA, translationally tagged with the mCherry fluorescent protein, from its native *hupA* locus, in addition to the chromosomally encoded Venus-CAT resistance protein. The response of this strain to Cp exposure at the MIC (30 µg/mL), as observed using the MCMA device, was consistent with that of F3-VCAT/pmCh; Cp exposure suppressed the growth and division of most cells, eventually leading to complete growth arrest, but a subpopulation of cells continued to grow and divide, ultimately passing stable resistance to their progeny (Fig. 6A–D and Movie S7).

**Figure 6:**
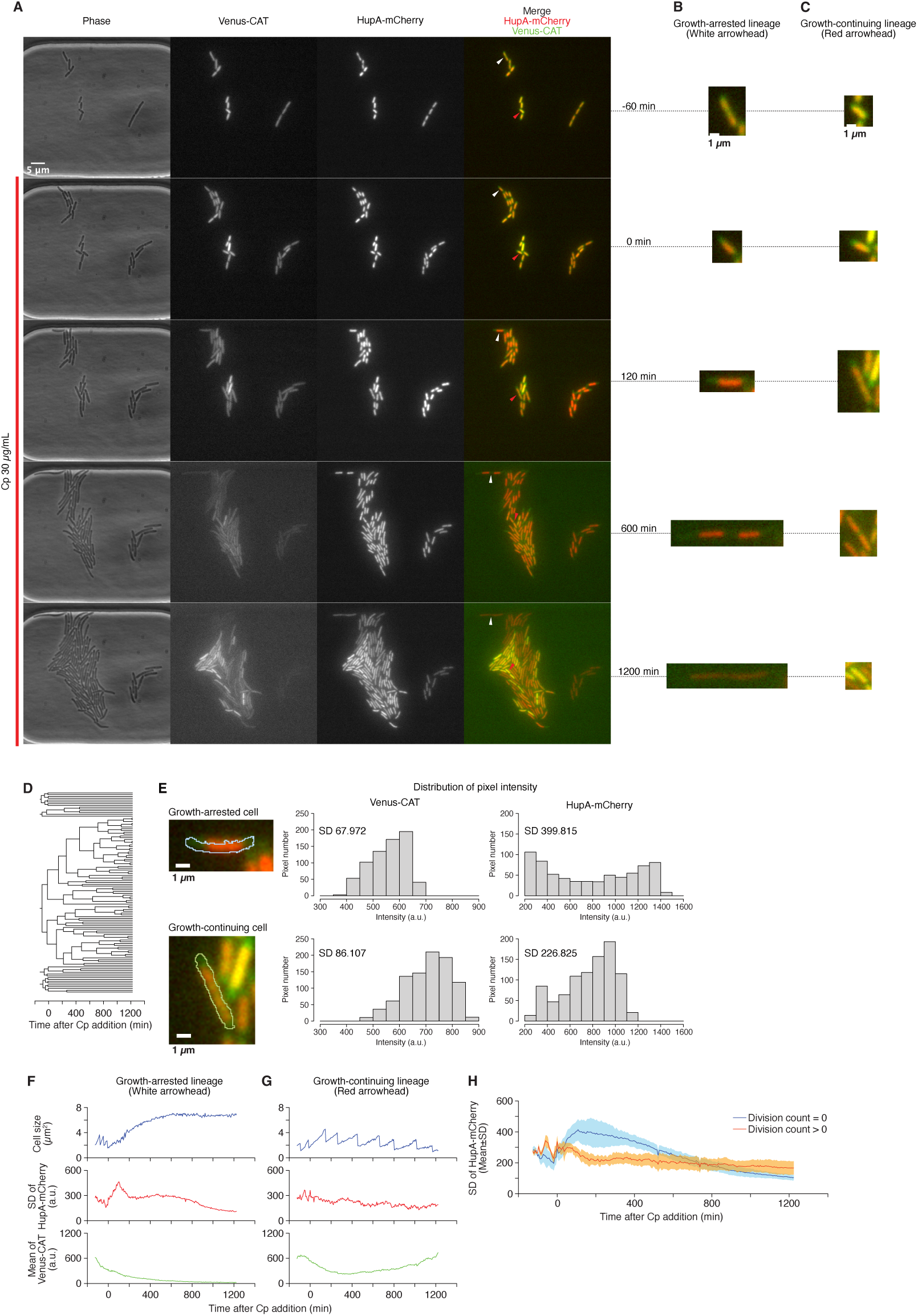
Spatial distribution of nucleoid in Cp-stressed cells. **A**. Time-lapse images of representative F3-VCAT-HupAmCh cells exposed to 30 µg/mL Cp in the MCMA device. The images show, from left to right: phase-contrast, Venus-CAT fluorescence, HupA-mCherry fluorescence, and merged images of Venus-CAT (green) and HupA-mCherry (red) channels. The red line to the left of the images indicates the period of Cp exposure. **B, C**. Enlarged micrographs of a representative growth-arrested cell (B, indicated by the white arrowhead in panel A) and a representative growth-continuing cell (C, indicated by the red arrowhead in panel A). **D**. Single-cell lineage tree of the microcolony shown in panel A. **E**. Quantification of nucleoid compaction. The left images show growth-arrested and growth-continuing F3-VCAT-HupAmCh cells at 120 min after the onset of 30 µg/mL Cp exposure (enlarged micrographs of those shown in panels B and C with outlines of the cell area). The histograms show the distributions of fluorescence signal intensity within the cell areas for Venus-CAT (middle) and HupA-mCherry (right). Nucleoid compaction causes a pronounced bimodal distribution of the HupA-mCherry signal, as shown in the growth-arrested cell, leading to an increase in the SD of the distribution. **F, G**. Time serious of cell size, standard deviation (SD) of HupA-mCherry, and mean fluorescence of Venus-CAT in the growth-arrested lineage (F, corresponding to panel B), and the growth-continuing lineage (G, corresponding to panel C). **H**. Time series of the SD of HupA-mCherry signals in immediately growth-arrested lineages (blue, no cell division under Cp exposure, *n* = 20) and other cell lineages (orange, at least one cell division under Cp exposure, *n* = 92). The bold lines and shaded areas represent mean values and SDs, respectively.

The time-lapse observations revealed that nucleoids visualized by HupA-mCherry became spatially compacted at the center of immediately growth-arrested cells in response to Cp exposure, whereas they remained broadly distributed in growth-continuing cells (Fig. 6B and C, Movie S7).

To quantitatively evaluate the spatial distribution of nucleoids inside cells, we calculated the standard deviation (SD) of HupA-mCherry fluorescence intensity across all pixels within each individual cell area. Because the fluorescence intensity distribution becomes bimodal when fluorescence signals are spatially restricted inside cells (Fig. 6E), a high SD of fluorescence intensity indicates nucleoid compaction, whereas a low SD indicates a relaxed structure. In cells that arrested growth immediately after Cp exposure, the SD of HupA-mCherry fluorescence transiently increased, indicating nucleoid compaction (Fig. 6B, F, and H). In contrast, SD values remained relatively stable in cells that divided more than once (Fig. 6C, G, and H). These results demonstrate that avoiding initial nucleoid compaction is a key hallmark of growing cells upon Cp exposure.

## Discussion

Our study demonstrates that *E. coli* cells expressing an antibiotic resistance protein develop enhanced resistance within only a few generations of exposure to antibiotic. We reveal that, at the single-cell level, enhanced resistance is characterized by distinct, remodeled gene expression profiles. We show that changes in expression level of the resistance protein alone cannot explain the long-term dynamics of resistance enhancement. Moreover, depending on the specific cellular lineage that is selected, enhanced resistance may either be maintained stably after the antibiotic is removed, or may rapidly revert back to pre-enhancement levels. Our findings are thus consistent with a model in which enhanced resistance can occur by multiple routes, involving different subsets of genes, which can be stably maintained or reversed without genetic changes.

While our analysis is restricted to a model strain expressing an introduced resistance protein, this genetic configuration allowed us to thoroughly investigate the contribution of a single dominant resistance protein (Venus-CAT) to stable resistance enhancement under prolonged Cp exposure (Fig. 4C). Furthermore, it enabled us to identify the importance of gene expression heterogeneity in individual cells during the initial period of Cp exposure (Fig. 4D). Crucially, these results uncovered a dynamic shift in the importance of resistance protein expression, plausibly driven by multi-generational physiological changes in cells (Fig. 4D). The distinct transcriptomic profiles observed across different resistance-enhanced cell lineages (Fig. 5, S7B, S8, and S10) indicate that there are multiple avenues to counteract Cp exposure. Although the mechanisms underlying the enhanced resistance in each lineage remain to be fully elucidated, the availability of multiple resistance states—and the phenotypic plasticity that enables individual cells within a population to transition to one of these states—may partially explain the high adaptability of antibiotic-exposed cells.

It also remains to be determined what causes a specific transcriptomic state in each cell lineage. Pre-existing phenotypic heterogeneity before antibiotic exposure, recently characterized by bacterial single-cell RNA sequencing [42], might have contributed to the diversification of resulting transcriptomic states observed in this study. Notably, our analysis revealed a correlation between the maintenance of relaxed nucleoid structures upon Cp exposure and growth continuation (Fig. 6). According to the ”transertion” hypothesis, a relaxed nucleoid structure is associated with active transcription and translation [43]. Therefore, maintaining active genome-wide transcription may allow an efficient search for higher resistance states. Understanding the linkage between nucleoid structures and the intrinsic exploratory dynamics proposed by several studies [11, 44, 45] will be an important topic for future research.

Although whole-genome sequencing detected no mutations in the resistance-enhanced cell populations (Table S1), we cannot exclude the possibility that mutations—such as those in repetitive or structurally complex genomic regions not fully resolved by short-read MiSeq sequencing—were present. Indeed, in unstable heteroresistance—a phenomenon in which a bacterial subpopulation exhibits transiently elevated resistance (typically defined as an 8-fold or greater increase in MIC) relative to the main population—a common mechanism is the occurrence of genetically unstable tandem amplifications of resistance genes [26, 27, 46]. Tandem gene amplifications occur at high frequencies in bacterial genomes, ranging between 10*^−^*^6^ and 10*^−^*^3^ per cell [47]. Furthermore, amplified resistance genes often impose a significant fitness cost, causing the population to revert to a susceptible state through selection within a limited number of generations (*<* 50 generations) in the absence of antibiotics [27]. However, the relatively high frequency of cells giving rise to resistance-enhanced lineages (12%) as well as the rapid loss of resistance in their progeny within short timeframes (8–9 generations, Movie S6) observed in our microfluidic time-lapse analysis suggest that genetic changes alone cannot account for all of the resistance enhancement phenomena observed in our study.

The initial cell fate diversification between growth arrest and continuation upon Cp exposure was previously observed in a similar *E. coli* model by Deris *et al.* [48]. In their study, this cell fate diversification was explained by transitions of individual cells to either a stable growth state or a growth-suspended state, driven by a mutual feedback between resistance protein expression and growth-correlated ribosomal abundance under translational inhibition conditions. The initial positive correlation between Venus-CAT expression and fitness observed in our study (Fig. 4D) is consistent with this model as a mean behavior (Fig. 2C and E). However, many individual cells reached a stably growing state without expressing the resistance protein at high levels. Furthermore, stable growth during the regrowing phase was enabled without high expression of the resistance protein (Fig. 4C). Therefore, our results indicate that understanding the single-cell dynamics and long-term behaviors under Cp exposure observed in this study requires an extended framework beyond this simple model.

The lineage-specific transcriptomic profiles of resistance-enhanced cell populations suggest that multiple resistance-enhancing mechanisms can operate within a single population. This heterogeneity in counteracting mechanisms across single-cell lineages may have clinical relevance, as secondary drug regimens assuming a single mechanism could fail to eradicate sub-populations that resist through alternative pathways. Therefore, elucidating the diversity of resistance enhancement mechanisms that emerge within clonal bacterial populations will have crucial implications for both basic and clinical microbiology.

## Methods

### *E. coli* strains and culture conditions

*E. coli* strains used in this study were derived from the W3110F3 strain, a derivative of the W3110 strain that carries a *∼*50-kbp genomic deletion encompassing 52 genes (approximately positions 274,372–324,783; Table S1), as well as knockouts of three genes (*flu*, *fimA*, and *fliC*) introduced to prevent cellular adhesion to internal surfaces of microfluidic devices. The W3110F3 strain is designated as “F3” in this study.

To confer resistance to Cp, the *venus* and chloramphenicol acetyltransferase (*cat*) genes were integrated at the *intC* locus on the chromosome to express the translationally fused Venus-CAT resistance protein under the control of the P_LtetO_*_−_*_1_ promoter [31]. Because the Tet repressor is absent, P_LtetO_*_−_*_1_ serves as a constitutive promoter in the strains used in this study. Strains carrying this *venus-cat* module are indicated by “-VCAT” in their names.

To visualize nucleoids in single cells, an *E. coli* strain expressing C-terminally fused HupA-mCherry protein from the native *hupA* locus on the chromosome was constructed. The strain carrying this *hupA-mcherry* module is indicated by “-HupAmCh” in the strain name.

Three plasmids—pmCh, pOE-RcsA, and pOE-CysB—were used in this study. All of these plasmids were constructed from the low-copy-number pMW118 plasmid, which carries the pSC101 replication origin (ori). The pmCh plasmid expresses the mCherry protein under the control of the P_LtetO_*_−_*_1_ promoter. The pOE-RcsA and pOE-CysB plasmids express RcsA or CysB, respectively, under the control of the IPTG-inducible P_LlacO_*_−_*_1_ promoter [31].

By combining these genetic elements, the following strains were constructed and used in this study: F3, F3-VCAT, F3/pmCh, F3-VCAT/pmCh, F3-VCAT/pOE-RcsA, F3-VCAT/pOE-CysB, and F3-VCAT-HupAmCh.

*E. coli* cells were cultured in M9 medium (1*×*M9 minimal medium (Difco) containing 2 mM MgSO_4_, 0.1 mM CaCl_2_, 0.2% glucose and 0.5*×*MEM amino acids solution (SIGMA)) at 37°C. Kanamycin (40 µg/mL) was added during overnight cultures for strains carrying pmCh, pOE-RcsA, or pOE-CysB.

### MIC measurement

A small amount of glycerol stock stored at -80°C was inoculated into 2 mL of M9 medium and cultured overnight at 37°C with shaking. Kanamycin (40 µg/mL) was added to overnight cultures of strains carrying pmCh, pOE-RcsA, or pOE-CysB. For resistance-enhanced cell populations collected from 96-well cultures, kanamycin was omitted to avoid its influence on cellular physiology, even though they carried the pmCh plasmid. To prepare exponentially growing cells, overnight cultures were diluted to an OD_600_ of 0.01 in 2 mL of fresh M9 medium and incubated at 37°C with shaking for 3-4 h. The cultures were then diluted to an OD_600_ of 0.0002. For assays using test tubes, 1 mL of the diluted culture was mixed with 1 mL of M9 medium containing Cp at 2*×* the indicated concentrations and incubated at 37°C with shaking for 23 h. For 96-well plate-based assays, 100 µL of diluted culture and 100 µL of M9 medium containing Cp at 2*×* the indicated concentrations were mixed in each well and incubated at 37°C for 23 h. The OD_600_ for each condition was measured after 23 h using a spectrophotometer (UV-1800, Shimadzu) or a plate reader (Epoch2, BioTek). The minimum inhibitory concentration (MIC) was defined as the lowest Cp concentration at which the OD_600_ was below 0.02 (spectrophotometer) or below 0.002 (plate reader).

### Microfluidic devices

Two types of microfluidic devices, the MCMA and the dynamics cytometer, were used in this study. These devices were fabricated as described previously [32, 33]. Briefly, microchambers in the MCMA device and channels in the dynamics cytometer were created on glass coverslips (NEO Micro glass, No. 1, 24 mm *×* 60 mm, Matsunami) by wet etching with a buffered hydrofluoric acid solution (110-BHF, Morita Kagaku Kogyo). The surface of the glass coverslips was chemically decorated with biotin to enclose *E. coli* cells in the microchambers or to capture cells in the growth channels using a streptavidin-decorated cellulose membrane cover. A pad made of polydimethylsiloxane (PDMS) was attached to the coverslip via a Frame-Seal Incubation Chamber (9 mm *×* 9 mm, 25 µL internal volume, Bio-Rad) to supply culture medium into the devices.

### Single-cell time-lapse observation

Overnight cultures were prepared as described in the MIC measurement section above and diluted 100-fold in 2 mL of fresh M9 medium, followed by incubation at 37°C with shaking for 3-4 h. Exponentially growing cells were loaded into microfluidic devices as described previously [32, 33]. The assembled microfluidic device was connected to silicone tubing to perfuse culture medium and placed on the microscope stage. A syringe pump was used to control the medium flow rate at 6 mL/h for the dynamics cytometer and 2 mL/h for the MCMA device. Temperature was maintained at 37°C using a microscope-mounted temperature control chamber (TIZHB; Tokai Hit) during time-lapse imaging. Time-lapse imaging was performed using a Nikon Ti-E microscope equipped with an ORCA-R2 camera or an ORCA-flash camera (Hama-matsu Photonics), a 100*×* objective lens (Plan Apo *λ*, 100*×*/NA1.45, oil immersion; Nikon), the temperature control chamber described above, and a computer running Micro-manager [49] for automated image acquisition. An LED illuminator (MCWHL2; Thorlabs) was used for fluorescence excitation. Phase contrast, Venus-CAT fluorescence, and mCherry fluorescence images were acquired.

### Image and data analysis

A custom Fiji [50] macro was used for cell segmentation and cell lineage tracking in time-lapse images. Errors in segmentation and lineage assignment were manually corrected. Numerical data specifying cell size (area), fluorescence intensity and its standard deviation (SD), and lineage relationships of individual cells were exported to text files for subsequent data analysis. The exported data were analyzed using a custom C program.

### Detection of non-dividing cell branches in single-cell pedigree trees

Single-cell time-lapse measurements with the MCMA device identified single cells that continued to grow and divide under Cp exposure as well as those that halted growth completely (Fig. 2, S1, and S2). The fates of cell branches in the single-cell pedigree trees–distinguishing between growth-continuing (surviving) and growth-arrested (dead)–were not evident for some of the branches at later timepoints in the measurements with the MCMA device. Therefore, the distinct fates of the branches reaching the defined end timepoint were estimated using a Bayesian inference approach that utilizes cellular traits.

Because the duration for which individual cells could be tracked varies among micro-colonies, we defined 1,701 min after Cp addition (630th frame) as the end timepoint of the analysis.

To characterize the traits of surviving and dead cells, information on four traits–mean Venus-CAT fluorescence intensity, mean mCherry fluorescence intensity, cell size, and age (time elapsed since the previous division)–was obtained from individual cells between 1,101 and 1,701 min (430th–630th frame). Based on this information, the probability distribution of each trait *X* conditional on fate, *P* (*X|*fate), was evaluated (Fig. S4B–E). The fates of individual cells at the defined end timepoint were inferred by sequentially updating the probability that the fate is dead, following Bayes’ theorem:

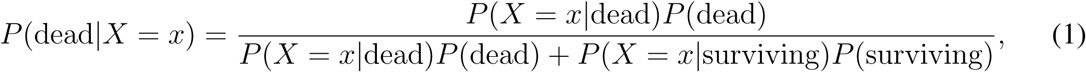

starting from an initial prior probability of *P* (dead) = *P* (surviving) = 1*/*2, where *X* is a trait and *x* is an observed trait value. The resulting *P* (dead*|X* = *x*) was used as a prior probability for the subsequent update with a different trait. When the final probability of a dead fate was larger than 0.5, we labeled the branch as ”dead”; otherwise, it was labeled as ”surviving.” The precision of the inference was verified using the data of microcolonies that were tracked beyond the defined end timepoint (Table S1).

To evaluate population dynamics under Cp exposure in the MCMA device, we excluded both observationally confirmed and inferred dead branches by truncating them immediately after their last division events (Fig. S4A, right).

### Limiting dilution culture

An exponentially growing cell culture of F3-VCAT/pmCh was prepared as described in the MIC measurement section above. A highly diluted cell suspension with an OD_600_ corresponding to 5.0*×*10^-9^ was prepared from the exponentially growing cell culture. For experiments to isolate resistance-enhanced cell lineages, the cell suspension contained 30 µg/mL Cp. This diluted cell suspension was dispensed at 200 µL per well into a 96-well plate. The plate was then sealed and placed in a plate reader (Epoch2; Biotek). The culture in each well was incubated at 37°C with shaking, and the OD_600_ was measured over time. Under this cell density condition, the number of cells per well was expected to be 0.78 (Fig. S5A).

### Microarray analysis

The growth of cells in the presence of 30 µg/mL Cp in each well prepared as a limiting dilution culture was monitored using a plate reader. Cells were collected from wells when the OD_600_ reached 0.3–0.6. A total of 150 µL of the collected culture was transferred to fresh M9 medium containing Cp (30 µg/mL) to a final volume of 2 mL and cultured again at 37°C until the OD_600_ reached approximately 0.4–0.5. For RNA extraction, 360 µL of the culture was mixed with a phenol–ethanol solution (0.1 g/mL), and cells were pelleted by centrifugation. The supernatant was removed, and the cell pellet was flash-frozen in liquid nitrogen and stored at -80°C until RNA extraction. In parallel, 750 µL of the culture was mixed with glycerol to a final concentration of 15% and stored at -80°C.

To obtain samples of the resistance-enhanced cell lineages under Cp-free conditions, the glycerol stocks prepared above were inoculated into 2 mL of M9 medium and incubated overnight at 37°C with shaking. As a control, cells from the original glycerol stock of F3-VCAT/pmCh were cultured in parallel. Kanamycin (40 µg/mL) was added only during the overnight culture of the original cells. Each overnight culture was diluted to an OD_600_ of 0.01 and incubated for 3–4 h. A portion of the culture was used to prepare a cell suspension at an OD_600_ of 2.0*×*10^-4^ for MIC measurements, while the remaining culture was grown to an OD_600_ of 0.4 for microarray analysis. For RNA extraction, 360 µL of the culture was mixed with a phenol–ethanol solution (0.1 g/mL), followed by centrifugation. The cell pellet was flash-frozen in liquid nitrogen and stored at -80°C. Another 750 µL portion was used to prepare glycerol stocks.

Total RNA was extracted using the RNeasy Micro Kit (QIAGEN). Subsequent microarray analysis was performed as described previously [8]. Quantile-nomalized gene expression data were used in Fig. 5A-C, S8, and S10. Genes absent from the genome were excluded from the plot in Fig. 5A-C, and S8.

### Whole-genome sequencing

Whole-genome sequencing was performed on seven samples: the A3, E9, and H10 cell lineages stocked immediately after cultivation under Cp exposure; those same three cell lineages stocked after cultivation under Cp-free conditions; and the original F3-VCAT/pmCh cell population. Cells from each glycerol stock were inoculated into M9 medium and cultured at 37°C with shaking. Once the OD_600_ exceeded 1.0, rifampicin was added to a final concentration of 0.3 mg/mL, and the cultures were incubated for an additional 3 h. Cells were then collected by centrifugation from 1 mL of culture, and the pellets were stored at -80°C until genomic DNA extraction.

Genomic DNA was extracted using the DNeasy Blood and Tissue Kit (QIAGEN). Subsequent sequencing and primary data processing were performed as described previously [8].

Genetic mutations in the samples were identified using breseq [37], using a custom reference genome constructed by integrating the *venus-cat* module into the *E. coli* W3110 genome (NCBI accession AP009048) alongside the deletion of the *fimA*, *flu*, and *fliC* genes, as well as including the sequence of the pmCh plasmid.

### India ink staining

A 5-µL aliquot of an *E. coli* culture was mixed with 5 µL of India ink on a coverslip. The mixture was gently overlaid with a second coverslip, and excess suspension was removed from the edges. The cell suspension with India ink was observed using a Nikon Ti-E microscope equipped with an ORCA-R2 camera (Hamamatsu Photonics) and a 100*×* objective lens (Plan Apo *λ*, 100*×*/NA1.45, oil immersion; Nikon). An LED illuminator (MCWHL2; Thorlabs) was used for fluorescence excitation. Images were acquired using Micro-manager [49].

### Materials availability

*E. coli* cell strains and vectors used in this study are available from the authors upon request.

## Acknowledgments

We thank Nozomu Obana for sharing the protocol of india ink staining; the members of the Wakamoto Lab for discussion. This work was supported by JST CREST Grant Number JP-MJCR1927 and JPMJCR26T4 (Y.W.); JST ERATO Grant Number JPMJER1902 (Y.W.); NIH Grant Number R01-GM097356 and R35-GM161539 (E.K.); Japan Society for the Promotion of Science KAKENHI Grant Number 17H06389, 19H03216, and 24H00552 (Y.W.); Shionogi Infectious Disease Research Promotion Foundation (Y.W.); and The Uehara Memorial Foundation (Y.W.).

## Supplemental figures

**Figure S1:**
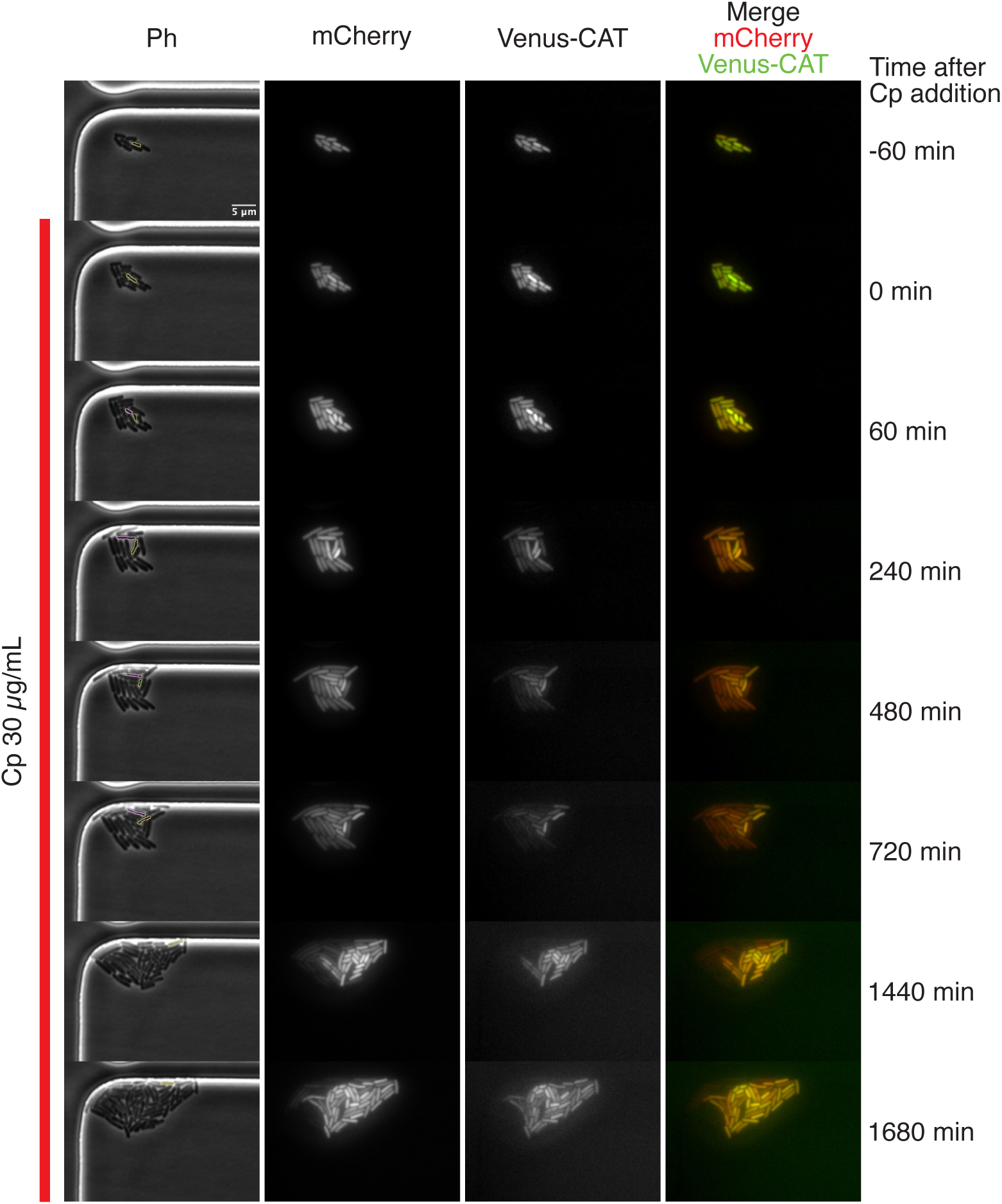
Additional example of the heterogeneous response of individual cells to Cp exposure. Time-lapse images show a representative F3-VCAT/pmCh microcolony exposed to 30 µg/mL Cp in the MCMA device. This microcolony is distinct from that shown in Fig. 2A. Displayed from left to right are phase-contrast, mCherry fluorescence, Venus-CAT fluorescence, and merged mCherry (red) and Venus-CAT (green) channels. The red line indicates the period of Cp exposure. A representative growing cell is outlined in yellow, and a representative growth-arrested cell is outlined in pink. These two cells are sister cells derived from the same cell division event after the onset of Cp exposure. The corresponding lineage tree and the transitions of cell size, mCherry fluorescent intensity, and Venus-CAT fluorescent intensity are shown in Fig. 2D and E.

**Figure S2:**
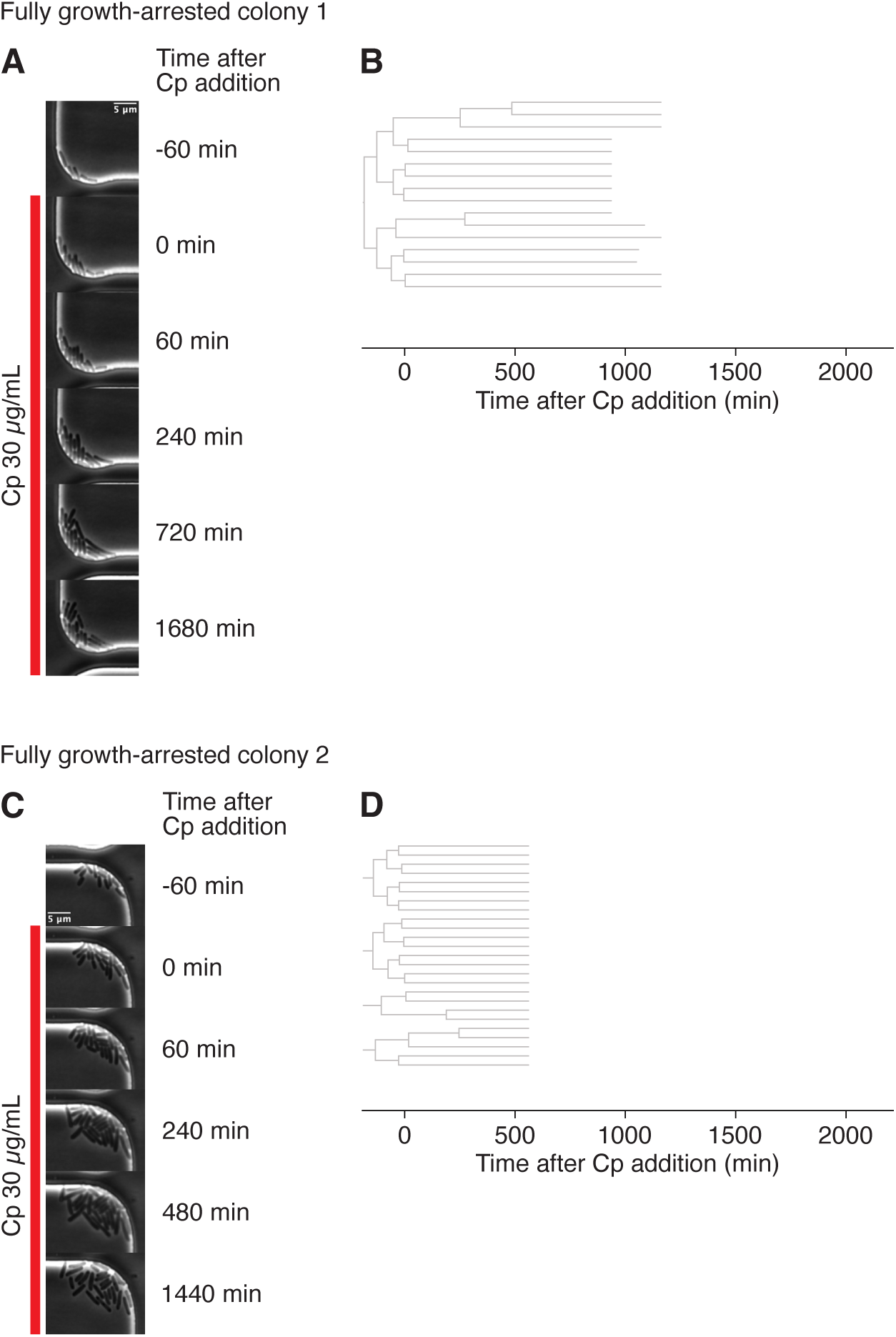
Representative fully growth-arrested colonies under Cp exposure. **A**. Time-lapse phase-contrast images of a representative F3-VCAT/pmCh microcolony exposed to 30 µg/mL Cp in the MCMA device, in which all cells stopped growing. The red line indicates the period of Cp exposure. Most cells exhibited cell body shrink-age following growth arrest (compare cell images at 720 min and 1680 min). **B**. Lineage tree corresponding to the microcolony shown in panel A. **C**. Time-lapse phase-contrast images of another representative F3-VCAT/pmCh microcolony exposed to 30 µg/mL Cp in the MCMA device that also underwent complete growth arrest. As in panel A, most cells exhibited cell body shrinkage following growth arrest (compare cell images at 480 min and 1440 min). **D**. Lineage tree corresponding to the microcolony shown in panel C.

**Figure S3:**
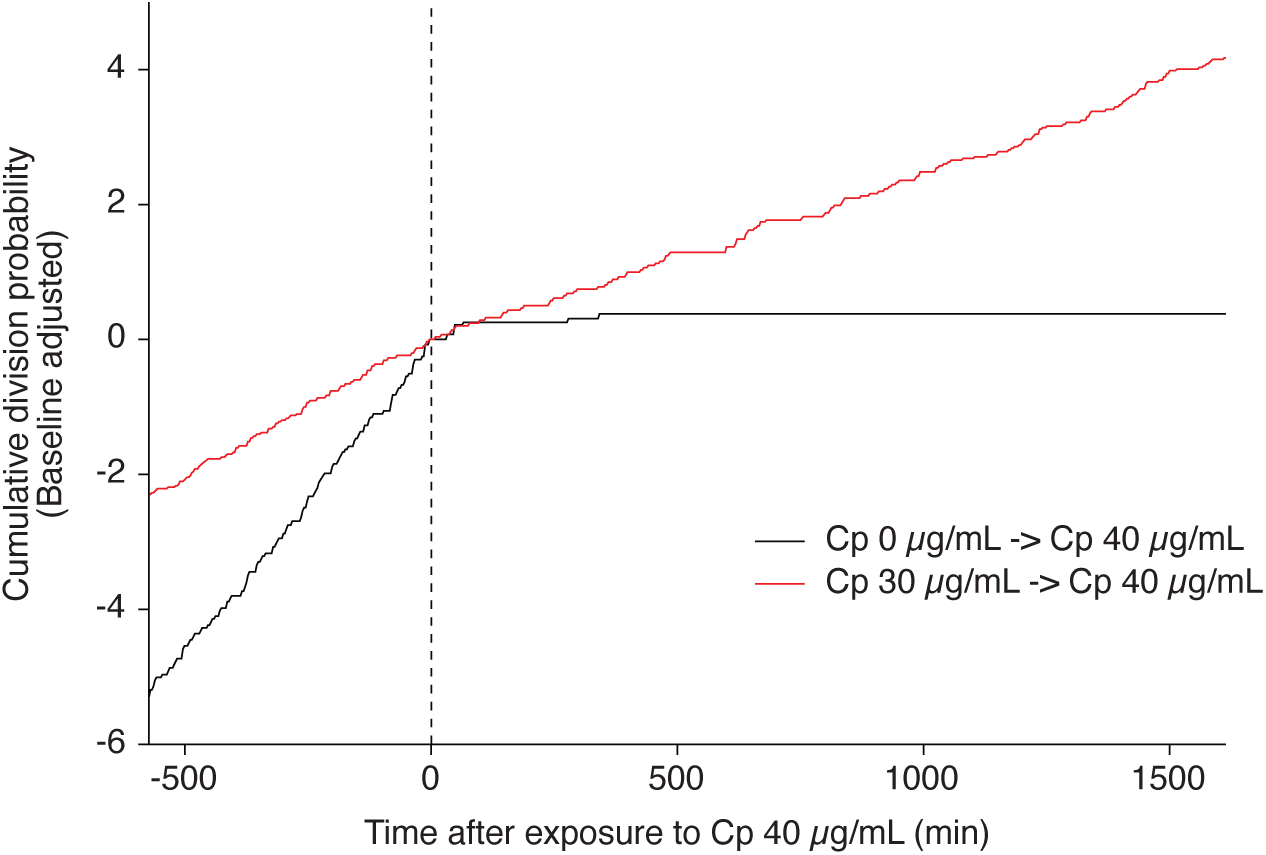
Verification of enhanced resistance of stably growing cells via a Cp concentration upshift. The division probability of cell populations in the dynamic cytometer was evaluated at each time point and summed over the time course as cumulative values. The black line represents cells initially cultured in Cp-free medium and subsequently exposed to 40 µg/mL Cp. The red line represents cells stably growing under exposure to 30 µg/mL Cp that were then exposed to 40 µg/mL Cp. Time zero (*t* = 0 min) indicates the onset of exposure to 40 µg/mL Cp. To compare population dynamics across conditions, the cumulative values at the time of the Cp concentration upshift were subtracted from the cumulative values.

**Figure S4:**
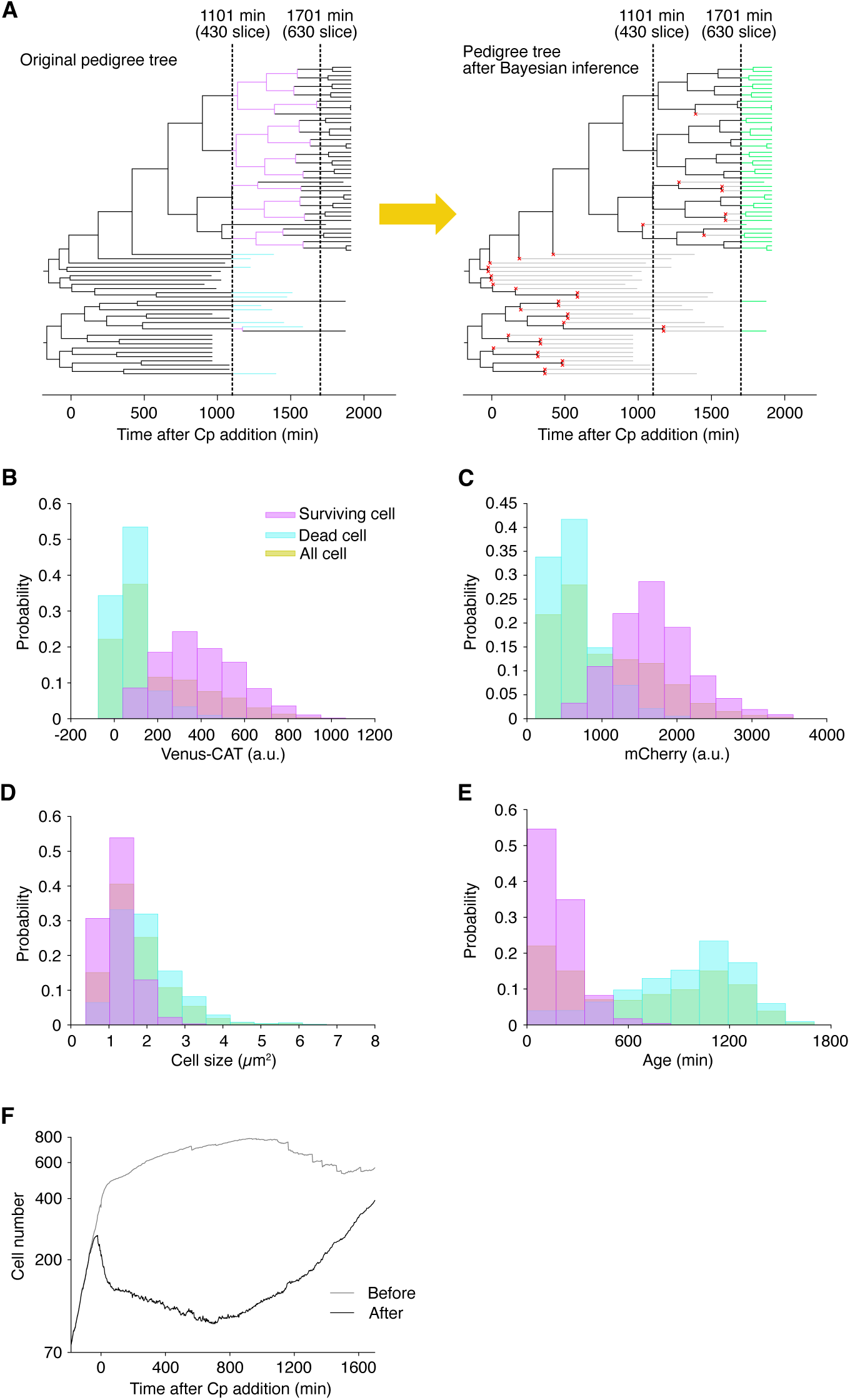
Detection of non-dividing cell branches in single-cell pedigree trees. **A**. Pedigree tree illustrating changes resulting from the removal of non-dividing cell branches. The left panel shows the original pedigree tree obtained by image analysis (the same as in Fig. 2B). Focusing on a 600-min time window from 1,101 min to 1,701 min (from the 430th to 630th time points in the time-lapse images), we identified “surviving (dividing) cell branches” (magenta) and “dead (non-dividing) cell branches” (cyan). Based on these branches, the distributions of four cellular traits–Venus-CAT fluorescence intensity, mCherry fluorescence intensity, cell size, and cell age (time elapsed since the previous division)–were evaluated for both surviving and dead branches. We thereby estimated the fate of each branch whose endpoint exceeds 1,701 min using Bayesian inference based on these four traits. The right panel shows the pedigree tree after the removal of dead cell branches and those inferred as dead. Branches with no further division events were truncated immediately after their last node (indicated by red crosses). Black branches represent viable cells, gray branches represent cells classified as dead, and green branches represent segments extending beyond 1,701 min. We defined 1,701 min (630th time point) as the endpoint for this analysis due to variable pedigree tree lengths across microcolonies. We confirmed that the false-positive rate of this inference method (incorrectly assigning a surviving cell as dead) was low (3.9%, 5/129; Table S1). **B–E**. Distributions of Venus-CAT fluorescence intensity (B), mCherry fluorescence intensity (C), cell size (D), and cell age (E) within the time window from 1,101 min to 1,701 min. Magenta boxes represent surviving cells, cyan boxes represent dead cells, and yellow boxes represent all cells. **F**. Changes in population dynamics resulting from the removal of dead cell branches in single-cell pedigree trees. The gray line indicates the total number of branches at each time point in the original pedigree (left panel of A), whereas the black line indicates the number of viable cell branches at each time point in the edited pedigree tree (black branches in the right panel of A). The population dynamics shown in black is identical to that presented in Fig. 4A.

**Figure S5:**
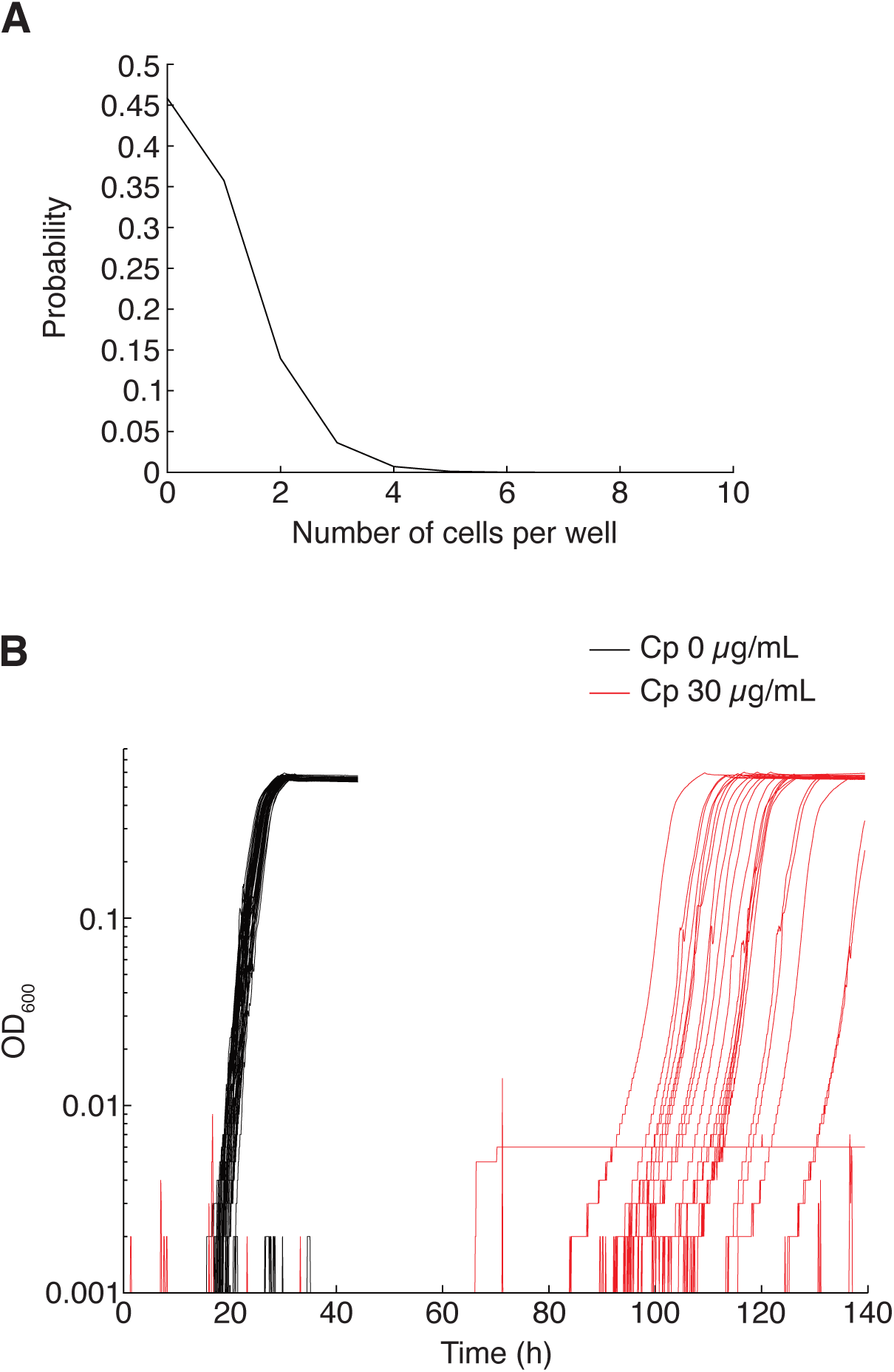
Limiting dilution cell cultures used in this study. **A**. Distribution of the number of cells per well, estimated using a Poisson distribution from the frequency of wells showing no growth under Cp-free conditions (44 out of 96 wells). The estimated mean number of cells per well was 0.78. **B**. Growth curves of F3-VCAT/pmCh cells observed in individual wells under the dilution conditions in A, either in Cp-free M9 medium (black) or in the presence of 30 µg/mL Cp (red). Cell growth was observed in 52 wells under Cp-free conditions (black lines) and in 22 wells under exposure to 30 µg/mL Cp (red lines).

**Figure S6:**
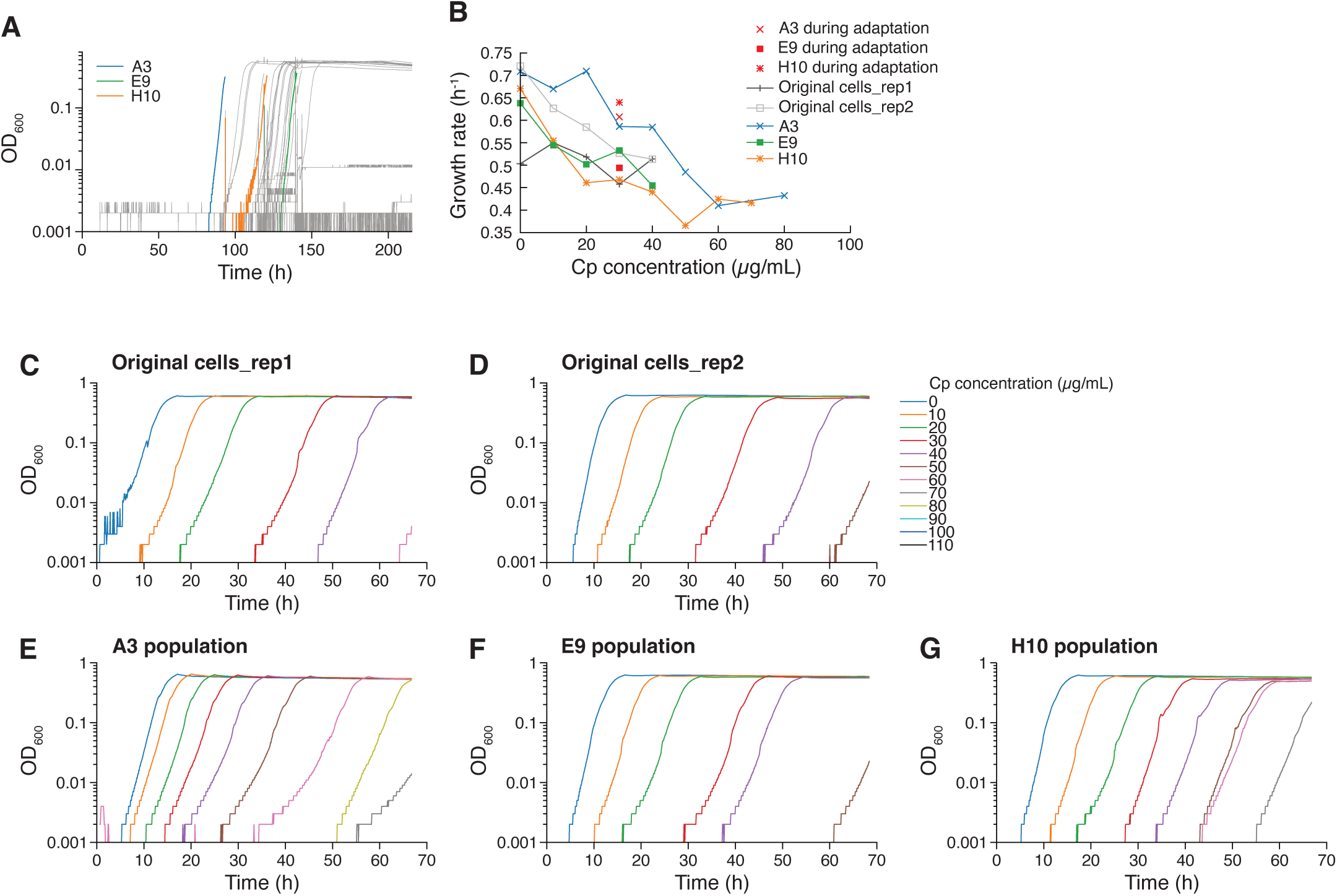
Growth of resistance-enhanced cell lineages in the presence or absence of Cp. **A**. Growth curves of cells obtained from limiting-dilution cultures in the presence of Cp. Each gray line represents a well in which growth was detected. Cells were sampled from a subset of these growth-positive wells, including the A3 (blue), E9 (green), and H10 (orange) cell lineages. The growth curves of the sampled cells are truncated at the time points of sampling. **B**. Growth rates of original cells and isolated resistance-enhanced A3, E9, and H10 cell lineages. Red symbols (cross for A3, filled square for E9, and asterisk for H10) indicate the growth rates of cultures in the presence of 30 µg/mL Cp in the limiting-dilution experiment shown in panel A. Other data points and lines represent growth rates measured during MIC determinations for each population (Fig. 5E). The corresponding growth curves are shown in panels C–G. The OD_600_ range of 0.04–0.2 was used to calculate each growth rate. **C–G**. Growth curves used for MIC determination against Cp of the original population (replicate 1, C; replicate 2, D), A3 (E), E9 (F), and H10 (G).

**Figure S7:**
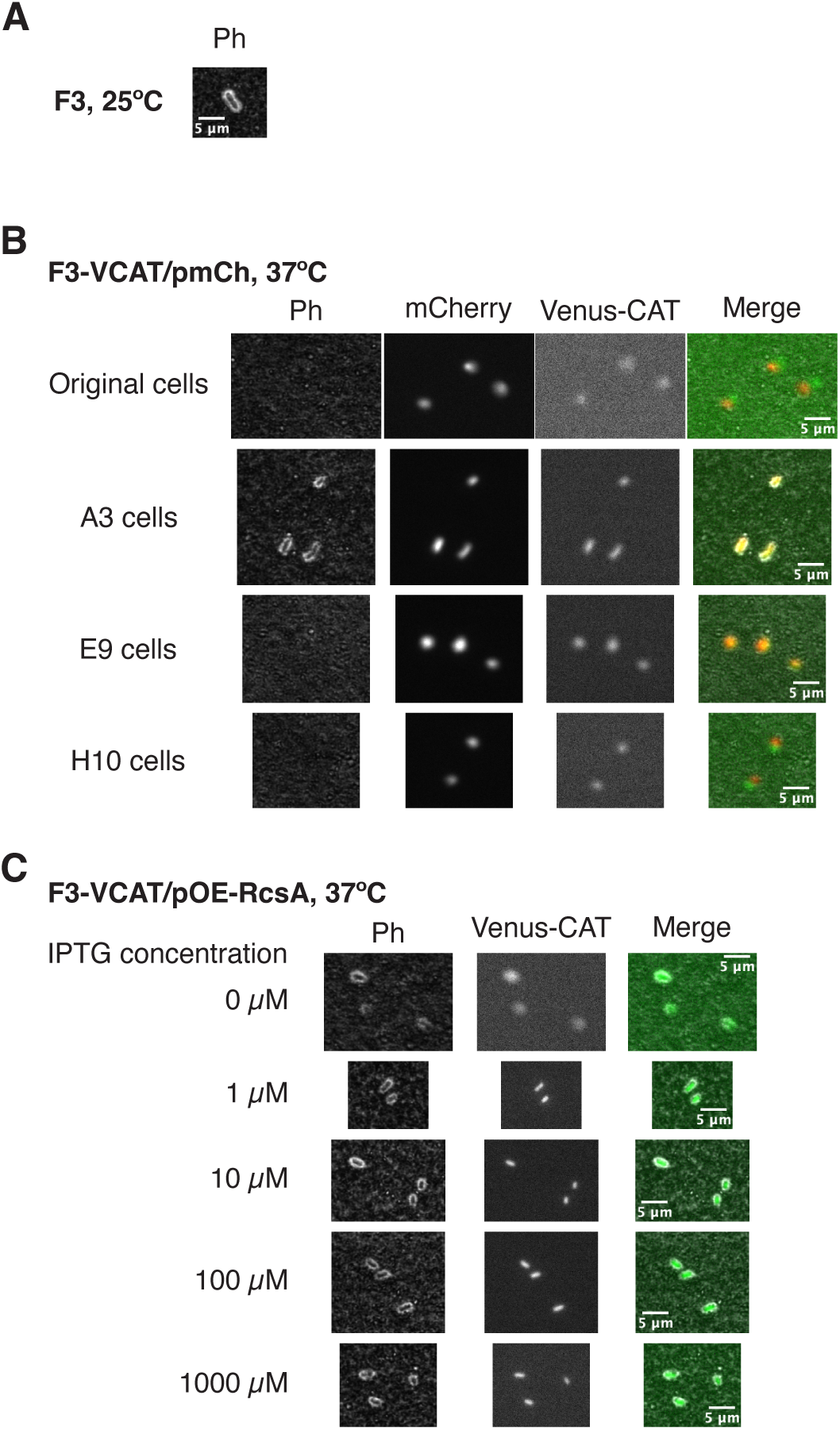
Capsule formation of cells at 37°C. **A**. India ink staining of F3 cells cultured at 25°C. A phase-contrast image is shown. The presence of a capsule can be detected as a void space around the cell into which the ink cannot penetrate. **B**. India ink staining of F3-VCAT/pmCh-derived cells (original, A3, E9, and H10) cultured at 37°C. From left to right, the images show phase-contrast, mCherry, Venus-CAT channels, followed by merged images of phase-contrast, mCherry, and Venus-CAT. Although all cells are surrounded by deep black India ink, only A3 cells (second row) are visible in the phase-contrast image, whereas the other cells, including original cells (top row), are hardly visible. **C**. India ink staining of F3-VCAT/pOE-RcsA cells cultured at 37°C induced with 0, 1, 10, 100, or 1000 µM IPTG. The additional copy of *rcsA* on a low-copy plasmid is inducible by IPTG in this strain. From left to right, the images show phase-contrast, Venus-CAT channel, and merged images of phase-contrast, and Venus-CAT. Capsule formation is observed around cells even at low induction levels.

**Figure S8:**
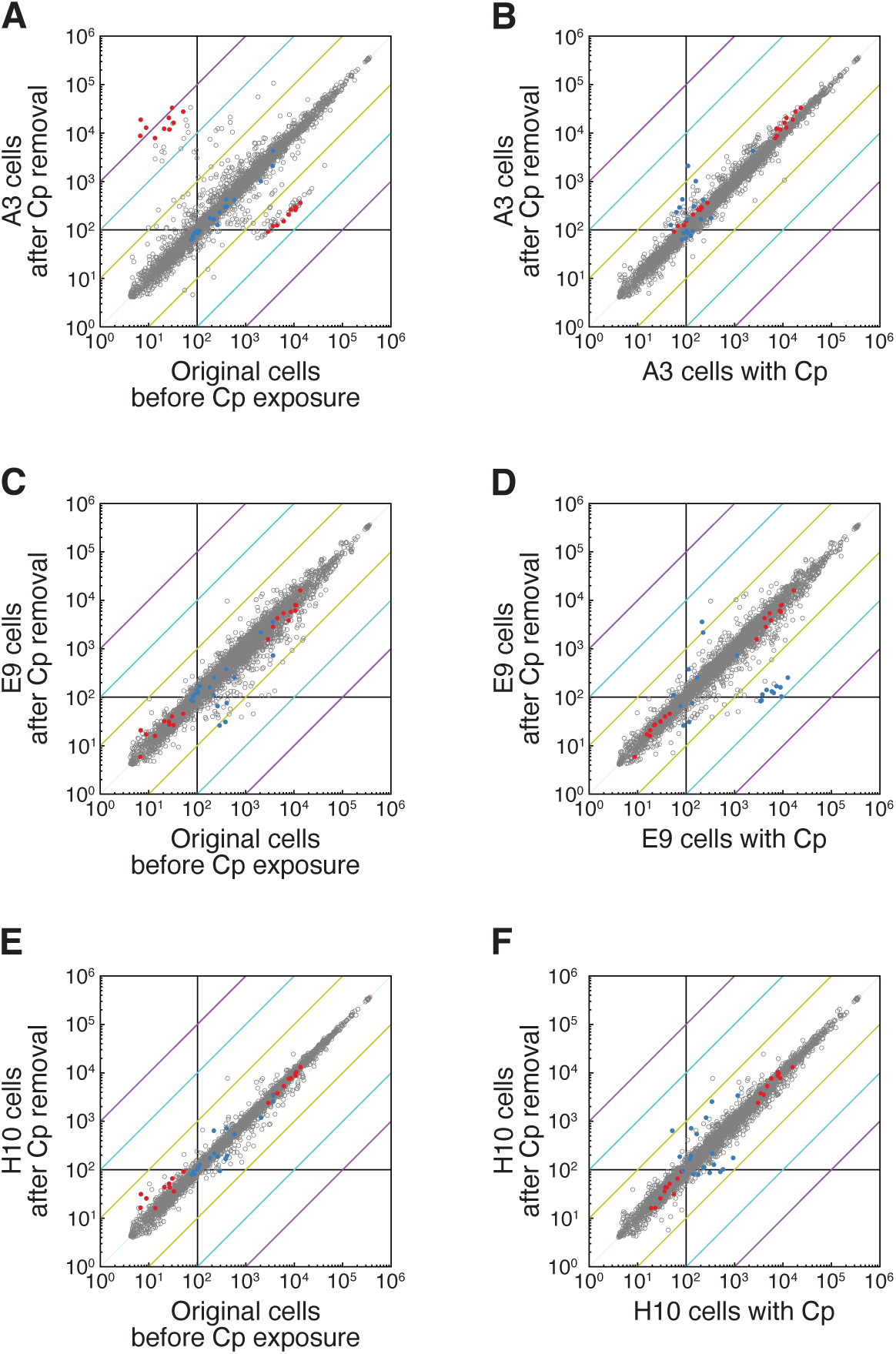
Genome-wide transcriptomic profiles of resistance-enhanced cell populations. **A–F**. Scatter plots illustrating global gene expression patterns. Panels A, C, and E compare the transcriptomic profiles of A3 (A), E9 (C), and H10 (E) cells under Cp-free conditions following regrowth in the presence of 30 µg/mL Cp (vertical axis) with that of unexposed original cells (horizontal axis). Panels B, D, and F compare the transcriptomic profiles of A3 (B), E9 (D), and H10 (F) cells under Cp-free conditions following regrowth in the presence of 30 µg/mL Cp (vertical axis) with their profiles under preceding Cp exposure (horizontal axis). Each dot corresponds to a single gene. Genes exhibiting the top 10 and bottom 10 fold changes in expressionin A3 and E9 under Cp treatment relative to the original cells before Cp exposure are highlighted in red and blue, respectively. Black lines represent the baseline threshold below which expression levels are considered too low for reliable interpretation. Yellow, cyan, and purple diagonal lines indicate 10-fold, 100-fold, and 1,000-fold changes, respectively, providing visual references for the magnitude of expression shifts.

**Figure S9:**
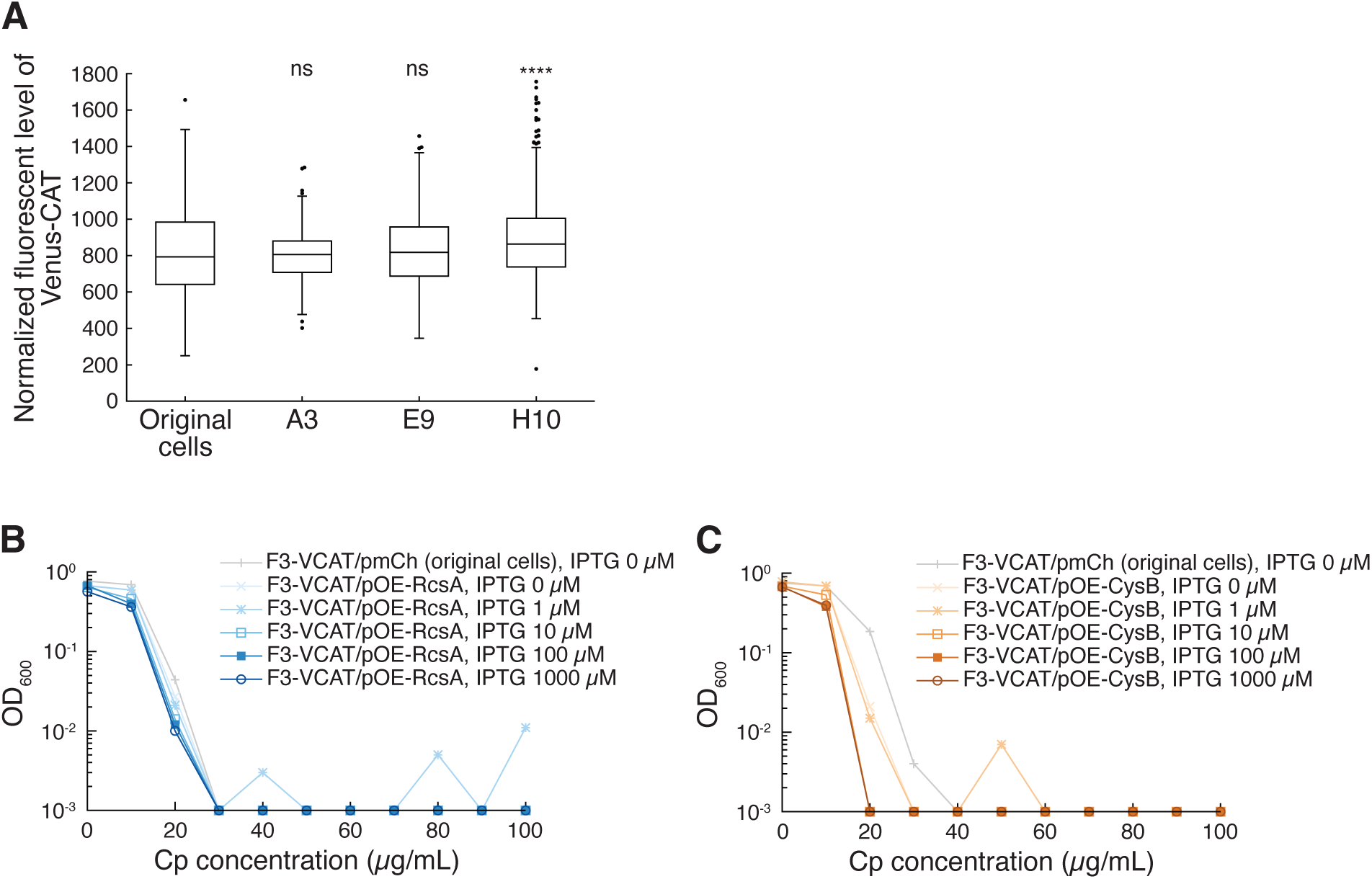
Contributions of Venus-CAT, RcsA, and CysB to Cp resistance. **A**. Comparison of Venus-CAT fluorescence levels among original, A3, E9, and H10 cells under Cp-free conditions. The distribution of Venus-CAT fluorescence levels in each population is shown as a box plot. The number of analyzed cells is 222 for original cells, 179 for A3, 369 for E9, and 650 for H10. Fluorescence signals were normalized across cell populations using the mean background fluorescence intensity. One-way ANOVA followed by Dunnett’s multiple comparisons test was used to detect differences from original cells (****, *p <* 0.0001; ns, *p >* 0.05). **B**. MIC measurements with RcsA overexpression. The F3-VCAT/pOE-RcsA strain, in which expression of an additional *rcsA* gene on the plasmid is inducible with IPTG, was used. The optical density at 600 nm of *E. coli* cell cultures after 23-h incubation at 37°C in M9 medium containing different concentrations of Cp (horizontal axis) is shown. Blue lines represent the results for the F3-VCAT/pOE-RcsA strain, with increasing color intensity corresponding to IPTG concentrations from 0 to 1,000 µM. The light gray line indicates the result for the F3-VCAT/pmCh strain without IPTG, used as a control. **C**. MIC measurements with CysB overexpression. The F3-VCAT/pOE-CysB strain, in which expression of an additional *cysB* gene on the plasmid is inducible with IPTG, was used. Orange lines represent the results for the F3-VCAT/pOE-CysB strain, with increasing color intensity corresponding to IPTG concentrations from 0 to 1,000 µM.

**Figure S10:**
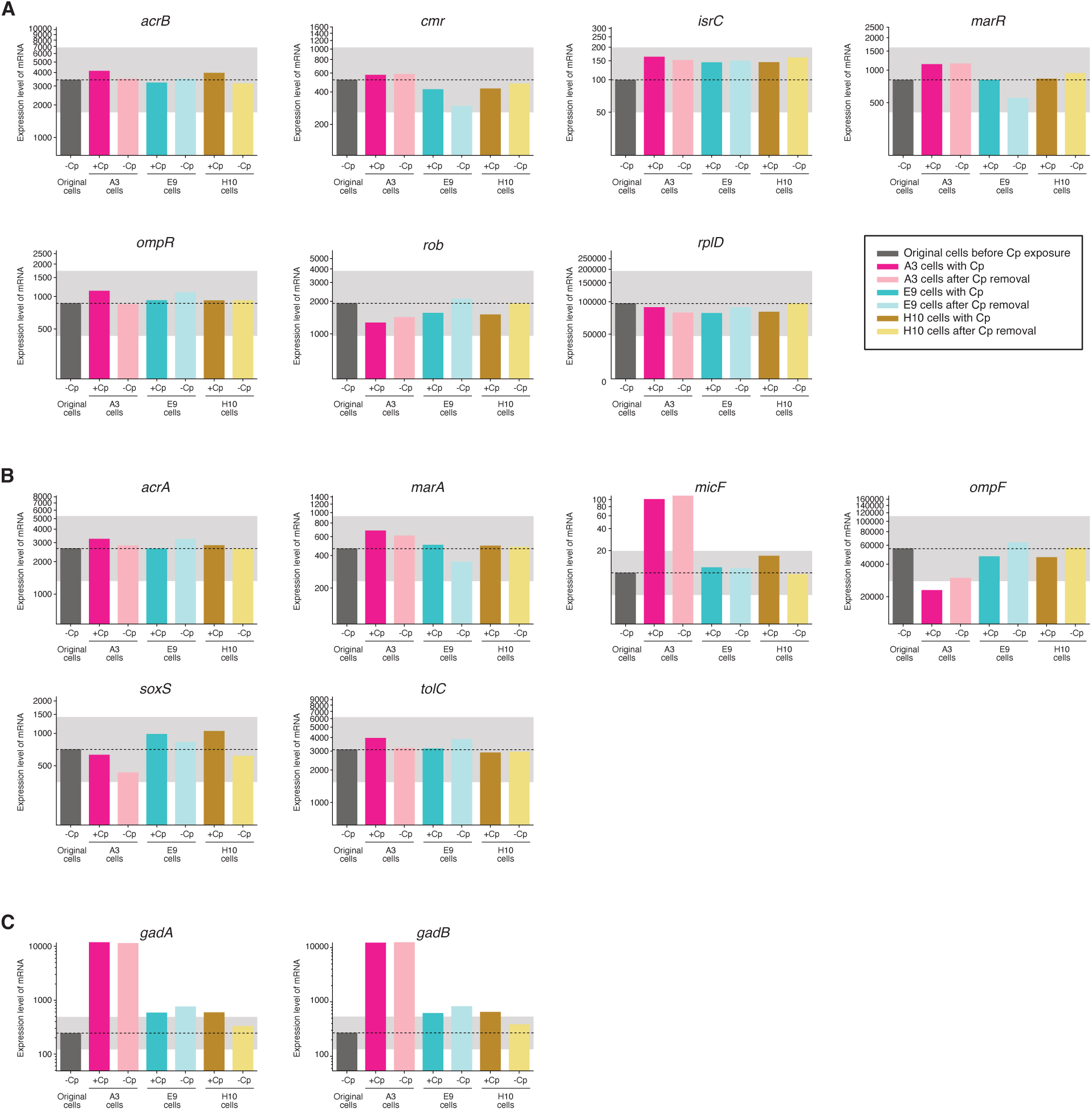
Expression levels of selected genes potentially underlying Cp resistance in the isolated resistance-enhanced cell lineages. **A**. Genes that acquired genetic alterations in a Cp-resistance evolution experiment in [6] (*acrB*, *cmr*, *isrC*, *marR*, *ompR*, *rob*, and *rplD*). **B**. Multidrug resistance genes (*acrA*, *marA*, *micF*, *ompF*, *soxS*, and *tolC*). **C**. Acid resistance genes (*gadA* and *gadB*). The gray shaded area in each plot represents a two-fold change range in gene expression relative to the unexposed parental cells.

## Supplemental tables

**Table S1:**
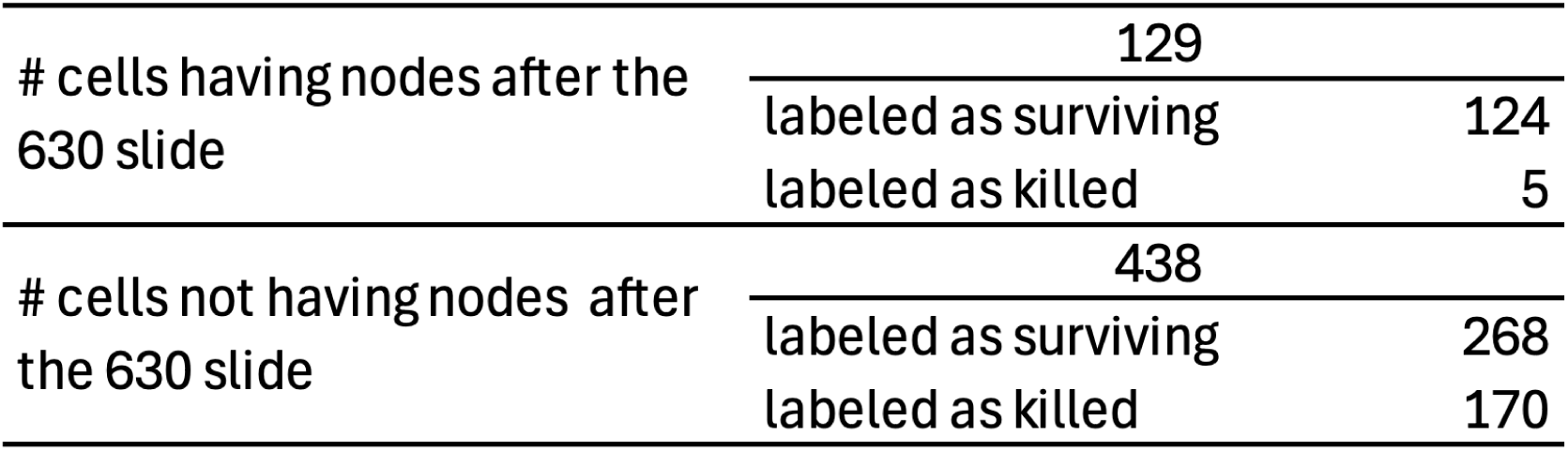
Summary of inferred fates of single-cell branches. This table summarizes the results of Bayesian estimation for the fates of single-cell branches that reached or extended beyond the defined end timepoint, 1,701 min (630th frame). A total of 567 single-cell branches reached the end timepoint in our dataset. These branches are categorized into two groups based on whether subsequent cell division was confirmed after the end timepoint. For each group, the numbers of branches classified as “surviving” and “dead” are shown. Because branches with confirmed cell divisions were unequivocally alive at the defined end timepoint, the fraction of branches labeled as “dead” in this group provides an estimate of the false-positive rate of this inference method (incorrectly classifying a surviving cell as “dead”).

**Table S2:**
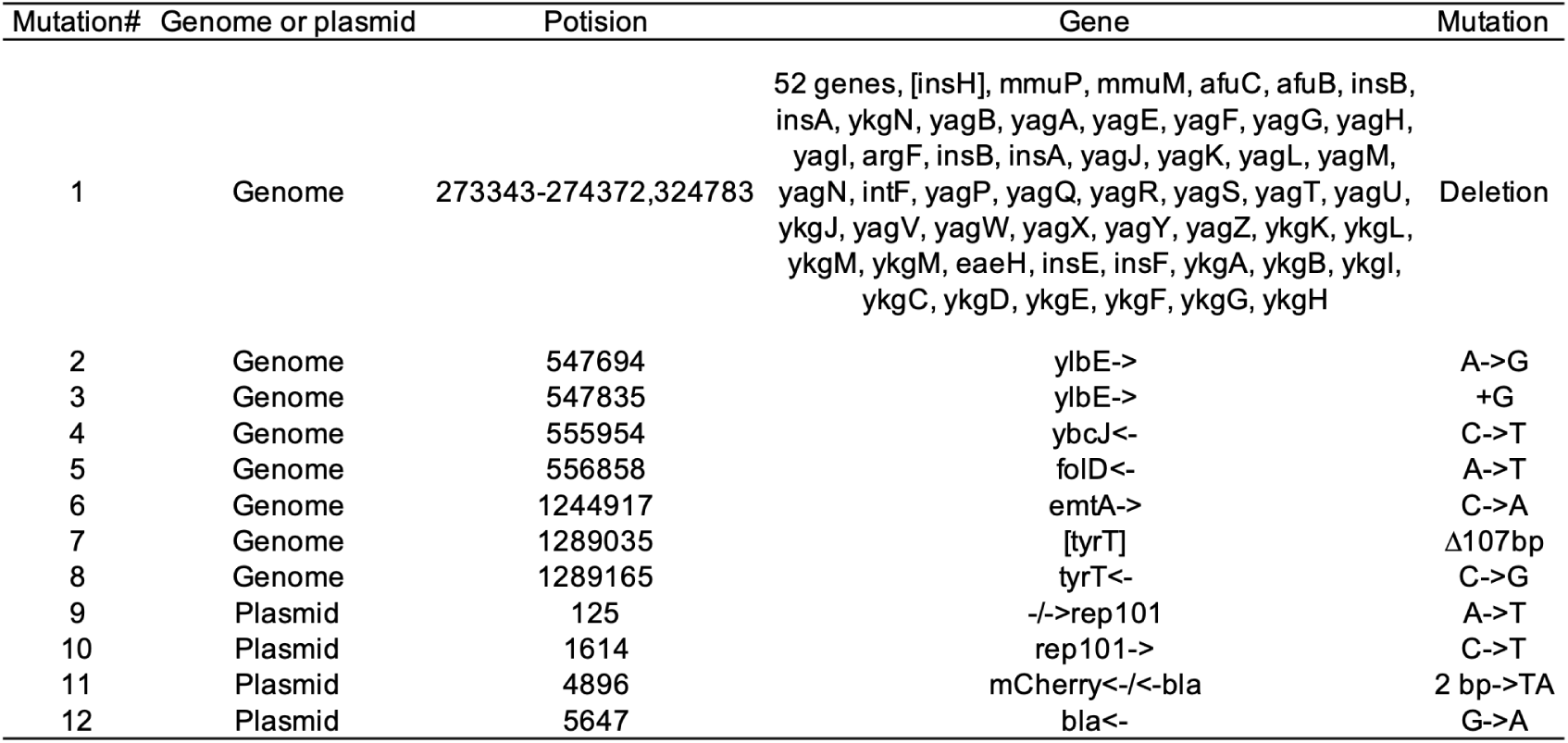
Genetic differences between the original F3-VCAT/pmCh cells and the reference genome. The genetic differences listed in this table were detected in the original F3-VCAT/pmCh cells. Identical differences were also found in the resistance-enhanced A3, E9, and H10 cell populations, but no additional genetic differences were detected in these cell lineages.

| Mutation# | Genome or plasmid | Potision | Gene | Mutation |
|---|---|---|---|---|
| 1 | Genome | 273343-274372,324783 | 52 genes, [insH], mmuP, mmuM, afuC, afuB, insB, insA, ykgN, yagB, yagA, yagE, yagF, yagG, yagH, yagI, argF, insB, insA, yagJ, yagK, yagL, yagM, yagN, intF, yagP, yagQ, yagR, yagS, yagT, yagU, ykgJ, yagV, yagW, yagX, yagY, yagZ, ykgK, ykgL, ykgM, ykgM, eaeH, insE, insF, ykgA, ykgB, ykgI, ykgC, ykgD, ykgE, ykgF, ykgG, ykgH | Deletion |
| 2 | Genome | 547694 | ylbE-> | A->G |
| 3 | Genome | 547835 | ylbE-> | +G |
| 4 | Genome | 555954 | ybcJ<- | C->T |
| 5 | Genome | 556858 | folD<- | A->T |
| 6 | Genome | 1244917 | emtA-> | C->A |
| 7 | Genome | 1289035 | [tyrT] | Δ107bp |
| 8 | Genome | 1289165 | tyrT<- | C->G |
| 9 | Plasmid | 125 | -/->rep101 | A->T |
| 10 | Plasmid | 1614 | rep101-> | C->T |
| 11 | Plasmid | 4896 | mCherry<-/<-bla | 2 bp->TA |
| 12 | Plasmid | 5647 | bla<- | G->A |

## Supplemental movie legends

**Movie S1. Time-lapse observation of the F3-CAT/pmCh microcolony shown in (Fig. 2A-C) under a step change in Cp concentration (0*→*30 µg/mL)** F3-VCAT/pmCh cells were first cultured in Cp-free M9 medium, then exposed to 30 µg/mL Cp in the MCMA. The images show, from left to right: phase-contrast, mCherry fluorescence, Venus-CAT fluorescence, and merged images of mCherry (red) and Venus-CAT (green) channels. Medium conditions and elapsed time after the onset of the Cp exposure are indicated at the lower-left corner. Scale bar, 5 µm.

**Movie S2. Time-lapse observation of the F3-CAT/pmCh microcolony shown in (Fig. 2D, E, and Fig. S1) under a step change in Cp concentration (0*→*30 µg/mL)** F3-VCAT/pmCh cells were first cultured in Cp-free M9 medium, then exposed to 30 µg/mL Cp in the MCMA. The images show, from left to right: phase-contrast, mCherry fluorescence, Venus-CAT flu-orescence, and merged images of mCherry (red) and Venus-CAT (green) channels. Medium conditions and elapsed time after the onset of the Cp exposure are indicated at the lower-left corner. Scale bar, 5 µm.

**Movie S3. Time-lapse observation of F3-VCAT/pmCh cells under a step change in Cp concentration (0***→***30 µg/mL)** F3-VCAT/pmCh cells were first cultured in Cp-free M9 medium, then exposed to 30 µg/mL Cp in the DC. Left: Venus-CAT channel; Right: mCherry channel. Medium conditions and elapsed time after the onset of the Cp exposure are indicated at the lower-left corner. In this chamber, a single cell adapted, resumed growth and continued dividing for over 100 generations during Cp exposure. Scale bar, 5 µm.

**Movie S4. Comparison of F3-VCAT/pmCh cell behavior under different exposure histories to Cp (0***→***40 µg/mL vs. 30***→***40 µg/mL)** In the left movie, F3-VCAT/pmCh cells were cultured in Cp-free M9 medium and then exposed to 40 µg/mL Cp in the DC. In the right movie, cells had already been exposed to 30 µg/mL Cp for 5,787 min and had adapted to that environment, after which the Cp concentration was increased to 40 µg/mL. The mCherry channels are shown. The overall experimental condition (Cp0*→*Cp40 or Cp30*→*Cp40) is labeled in the upper left cormer of each movie. Midium conditions and the elapsed time after the onset of 40 µg/mL Cp exposure are indicated at the lower left. Scale bar, 5 µm.

**Movie S5. Time-lapse observation of F3-VCAT/pmCh cells under stepwise changes in Cp concentration (0***→***30***→***40***→***50***→***60 µg/mL)** F3-VCAT/pmCh cells were cultured in Cp-free M9 medium and then exposed to 30 µg/mL Cp for 5,787 min, 40 µg/mL Cp for 4,290 min, 50 µg/mL Cp for 4,290 min, and finally 60 µg/mL Cp in the DC. The mCherry channel is shown. Medium conditions are indicated at the upper left, and elapsed time after the onset of 30 µg/mL Cp exposure is shown at the lower left. In this experiment, the drain cahnnels eventually became clogged with cells, which terminated the observation. A portion of this movie is also shown in Movie S4 (right). Scale bar, 5 µm.

**Movie S6. Time-lapse observation of F3-VCAT/pmCh cells under stepwise changes in Cp concentration (0***→***30***→***0***→***30 µg/mL)** F3-VCAT/pmCh cells were cultured in Cp-free M9 medium, then exposed to 30 µg/mL Cp in the DC for 47.45 h, followed by 8.1 h in Cp-free medium, and subsequently re-exposed to 30 µg/mL Cp. Left: Venus-CAT channel; Right: mCherry channel. Medium conditions and elapsed time after the onset of the first Cp exposure are shown at the lower-left corner. In this chamber, cells adapted and resumed growth during the first Cp exposure, but failed to regrow during the second Cp exposure. Scale bar, 5 µm.

**Movie S7. Time-lapse observation of the F3-CAT-HupAmCh microcolony shown in (Fig. 6) under a step change in Cp concentration (0*→*30 µg/mL)** F3-VCAT-HupAmCh cells were first cultured in Cp-free M9 medium, then exposed to 30 µg/mL Cp in the MCMA. The images show, from left to right: phase-contrast, Venus-CAT fluorescence, HupA-mCherry fluorescence, and merged images of Venus-CAT (green) and HupA-mCherry (red) channels. Medium conditions and elapsed time after the onset of the Cp exposure are indicated at the lower-left corner. Scale bar, 5 µm.

